# Structural basis for the Rad6 activation by the Bre1 N-terminal domain

**DOI:** 10.1101/2022.10.23.513400

**Authors:** Meng Shi, Jiaqi Zhao, Mengfei Li, Wei Huang, Xue Bai, Wenxue Zhang, Kai Zhang, Xuefeng Chen, Song Xiang

## Abstract

The mono-ubiquitination of the histone protein H2B (H2Bub1) is a highly conserved histone post-translational modification that plays critical roles in many fundamental processes. In yeast, this modification is catalyzed by the conserved Bre1-Rad6 complex. Bre1 contains a unique N-terminal Rad6 binding domain (RBD), how it interacts with Rad6 and contributes to the H2Bub1 catalysis is unclear. Here, we present crystal structure of the Bre1 RBD-Rad6 complex and structure-guided functional studies. Our structure provides a detailed picture of the interaction between the dimeric Bre1 RBD and a single Rad6 molecule. We further found that the interaction stimulates Rad6’s enzymatic activity by allosterically increasing its active site accessibility and likely contribute to the H2Bub1 catalysis through additional mechanisms. In line with these important functions, we found that the interaction is crucial for multiple H2Bub1-regulated processes. Our study provides molecular insights into the H2Bub1 catalysis.

## Introduction

The basic unit of the eukaryotic chromatin, nucleosome core particle (NCP), is composed of a protein core consisting of two copies of histone proteins H2A, H2B, H3 and H4 and double strand DNA wrapped around it(Luger *et al*, 1997). Posttranslational modifications (PTMs) on the histone proteins play critical roles in regulating a variety of fundamental processes(Kouzarides, 2007; Strahl & Allis, 2000). Mono-ubiquitination of a conserved lysine residue in H2B (Lys123 in the budding yeast, Lys120 in mammals, H2Bub1) is a highly conserved histone PTM found in eukaryotes(West & Bonner, 1980). It is associated with actively transcribed genes, where it facilitates the gene transcription by loosing the chromatin(Fierz *et al*, 2011) and recruiting the FACT histone chaperon complex(Fleming *et al*, 2008; Pavri *et al*, 2006). It also regulates other histone protein PTMs associated with actively transcription, including H3K4 and H3K79 methylation catalyzed by DOT1L and the COMPASS complex, respectively(Briggs *et al*, 2002; Sun & Allis, 2002). Recent structural studies indicate that the ubiquitin molecule attached to H2B mediates interactions with DOT1L and the COMPASS complex, contributing to their recruitment and activation(Anderson *et al*, 2019; Hsu *et al*, 2019; Jang *et al*, 2019; Valencia-Sanchez *et al*, 2019; Worden *et al*, 2019; Worden *et al*, 2020; Xue *et al*, 2019; Yao *et al*, 2019). At DNA double strand breaks (DSBs), H2Bub1 facilitates the recruitment of factors for both homologous recombination (HR) and non-homologous end joining pathways, promoting DSB repair(Moyal *et al*, 2011; Nakamura *et al*, 2011; Shiloh *et al*, 2011; Zheng *et al*, 2018). During meiosis, H2Bub1 plays a critical role in the meiotic recombination required for the exchange of genetic materials between homologous chromosomes(Wang *et al*, 2017; Xu *et al*, 2016). H2Bub1 also has important functions in additional processes including DNA replication, nucleosome positioning, RNA processing, chromatin segregation and others(Fuchs & Oren, 2014). In line with its important cellular function, aberrant H2Bub1 levels in humans are implicated in several types of cancer(Marsh & Dickson, 2019; Marsh *et al*, 2020; Zhou *et al*, 2021).

The multiple-step ubiquitination reaction requires concerted actions of several enzymes. The ubiquitin activating enzyme (E1) conjugates ubiquitin to the ubiquitin-conjugating enzyme (E2), the ubiquitin ligase (E3) subsequently transfers ubiquitin from the E2 enzyme to the substrate. Whereas most organisms contain as few as one or two E1 enzymes and tens of E2 enzymes, they contain a few hundred E3 enzymes, which play critical roles in substrate selection(Deol *et al*, 2019; Stewart *et al*, 2016). In the budding yeast, the H2Bub1 reaction is catalyzed by the E3 enzyme Bre1 together with the E2 enzyme Rad6(Hwang *et al*, 2003; Robzyk *et al*, 2000; Wood *et al*, 2003). Bre1 is highly conserved among eukaryotes. The human Bre1 orthologs, RNF20 and RNF40, also cooperate with the human Rad6 orthologs (Rad6A and Rad6B) to catalyze the H2Bub1 formation(Kim *et al*, 2009; Kim *et al*, 2005; Zhu *et al*, 2005). Bre1, RNF20 and RNF40 belong to the RING domain of E3 enzymes and contain a C-terminal RING domain. The RING domain facilitates the ubiquitin transfer to the substrate by promoting the “closed” conformation of the E2~ubiquitin conjugate(Zheng & Shabek, 2017). Bre1’s RING domain also interacts with an acidic patch in NCP and positions Rad6 for the H2Bub1 catalysis(Gallego *et al*, 2016). In line these important functions, it was found that removing Bre1’s RING domain abolished H2Bub1 *in vivo*(Hwang *et al*., 2003), and the Bre1 fragment containing the RING domain and a predicted coiled-coil N-terminal to it was able to catalyze H2Bub1 *in vitro*(Turco *et al*, 2015).

In addition to the RING domain, Bre1 contains a N-terminal Rad6 binding domain (RBD) that also makes important contributions to the H2Bub1 catalysis(Kim & Roeder, 2009; Turco *et al*., 2015). However, the mechanism of its interaction with Rad6 and how it contributes to the H2Bub1 catalysis are poorly understood. Here, we present crystal structure of the Bre1 RBD-Rad6 complex and structure-guided functional experiments. Our structure revealed detailed mechanism of the Bre1 RBD-Rad6 interaction within the 2:1 Bre1 RBD-Rad6 complex. We found that the interaction stimulates Rad6’s enzymatic activity by allosterically increasing its active site accessibility and likely contribute to the H2Bub1 catalysis through additional mechanisms. In line with these important functions, we found that the interaction plays crucial roles in multiple H2Bub1-regulated processes inside the cell.

## Results

### Overall structure of the Bre1 RBD-Rad6 complex

The N-terminal 210 residues in the *Saccharomyces cerevisiae* Bre1 (ScBre1) have been reported to interact with Rad6 and contribute to the H2Bub1 catalysis(Kim & Roeder, 2009; Turco *et al*., 2015). This region is highly conserved among fungal Bre1 proteins (Fig. S1). We screened through several fungal species and were able to crystallize the *Kluyveromyces lactis* Rad6 (KlRad6) in complex with the *Kluyveromyces lactis* Bre1 (KlBre1) N-terminal fragment 1-206 or 1-184 (crystal forms 1 and 2, respectively). The Bre1 N-terminal region is predicted to contain several coiled-coils(Kim & Roeder, 2009; Turco *et al*., 2015). Using a predicted coiled-coil structure(Guzenko & Strelkov, 2018) and the structure of the *Saccharomyces cerevisiae* Rad6 (ScRad6, PDB 1AYZ)(Worthylake *et al*, 1998) as search models, we determined structures of crystal form 1 and 2 with molecular replacement. The structures were refined to resolutions of 3.2 Å and 3.05 Å (Table 1).

**Table 1.** Data collection and structure refinement statistics

|  | Crystal form 1 | Crystal form 2 |
| --- | --- | --- |
| Data collection |  |  |
| Space group | P6<sub>5</sub>22 | P6<sub>1</sub>22 |
| Cell dimensions |  |  |
| *a*, *b*, *c* (Å) | 94.44, 94.44, 534.64 | 113.22, 113.22, 386.17 |
| $\alpha$, $\beta$, $\gamma$ (°) | 90, 90, 120 | 90, 90, 120 |
| Resolution (Å) | 50-3.20 (3.26-3.2) | 50-3.05 (3.10-3.05) |
| R<sub>merge</sub> | 0.200 (1.797) | 0.092 (0.863) |
| I / σI | 17.0 (2.0) | 21.6 (1.1) |
| CC<sub>1/2</sub> | 0.985 (0.623) | 1.00 (0.876) |
| Completeness (%) | 100.0 (99.0) | 99.7 (96.9) |
| Redundancy | 18.0 (18.5) | 20.6 (16.7) |
| Refinement |  |  |
| Resolution (Å) | 40.89-3.20 (3.35-3.20) | 32.18-3.05 (3.11-3.05) |
| No. reflections | 22,676 (2,465) | 28,706 (2,495) |
| R<sub>work</sub> / R<sub>free</sub> (%) | 21.32/26.33<br>(32.57/39.34) | 23.72/28.60<br>(40.29/46.90) |
| No. atoms |  |  |
| Protein | 7563 | 7486 |
| Ligand/ion | 0 | 17 |
| B-factors |  |  |
| Protein | 107.63 | 122.41 |
| Ligand/ion | - | 118.27 |
| R.m.s. deviations |  |  |
| Bond lengths (Å) | 0.003 | 0.003 |
| Bond angles (°) | 0.599 | 0.645 |

\*Values in parentheses are for the highest-resolution shell.

The structures indicate that two KlBre1 N-terminal polypeptides dimerize to form an elongated RBD that binds to one KlRad6 molecule (Fig. 1A). In both crystal forms, residues 12-173 and 16-182 for KlBre1 polypeptides 1 and 2, respectively, are resolved in the electron density map. Additional KlBre1 residues included in the expression constructs are presumably disordered. Residues 1-158 in KlRad6 are resolved in the electron density map, part of its acidic C-terminal tail is disordered. Each KlBre1 polypeptide contains two coiled-coil regions (CC1 and CC2), which form extensive parallel coiled-coils interactions with the same regions in the other KlBre1 polypeptide. Each KlBre1 polypeptide also contains a domain M located between CC1 and CC2 that folds CC2 back to interact with CC1. Interactions between CC1 and CC2 and between these coiled-coils and domain M also contribute to the stabilization the RBD structure. The structures of CC1, CC2 and domain M in both KlBre1 polypeptides are quite similar. However, there are striking structural differences in the domain M-CC2 linker. With domain M or CC2 aligned, this linker in the two KlBre1 polypeptides point to opposite directions (Fig. 1B). There are also structural differences at the N- and C-terminus of the two KlBre1 polypeptides (Fig. 1B). As a result, the KlBre1 RBD dimer is strongly asymmetric.

**Figure 1.**
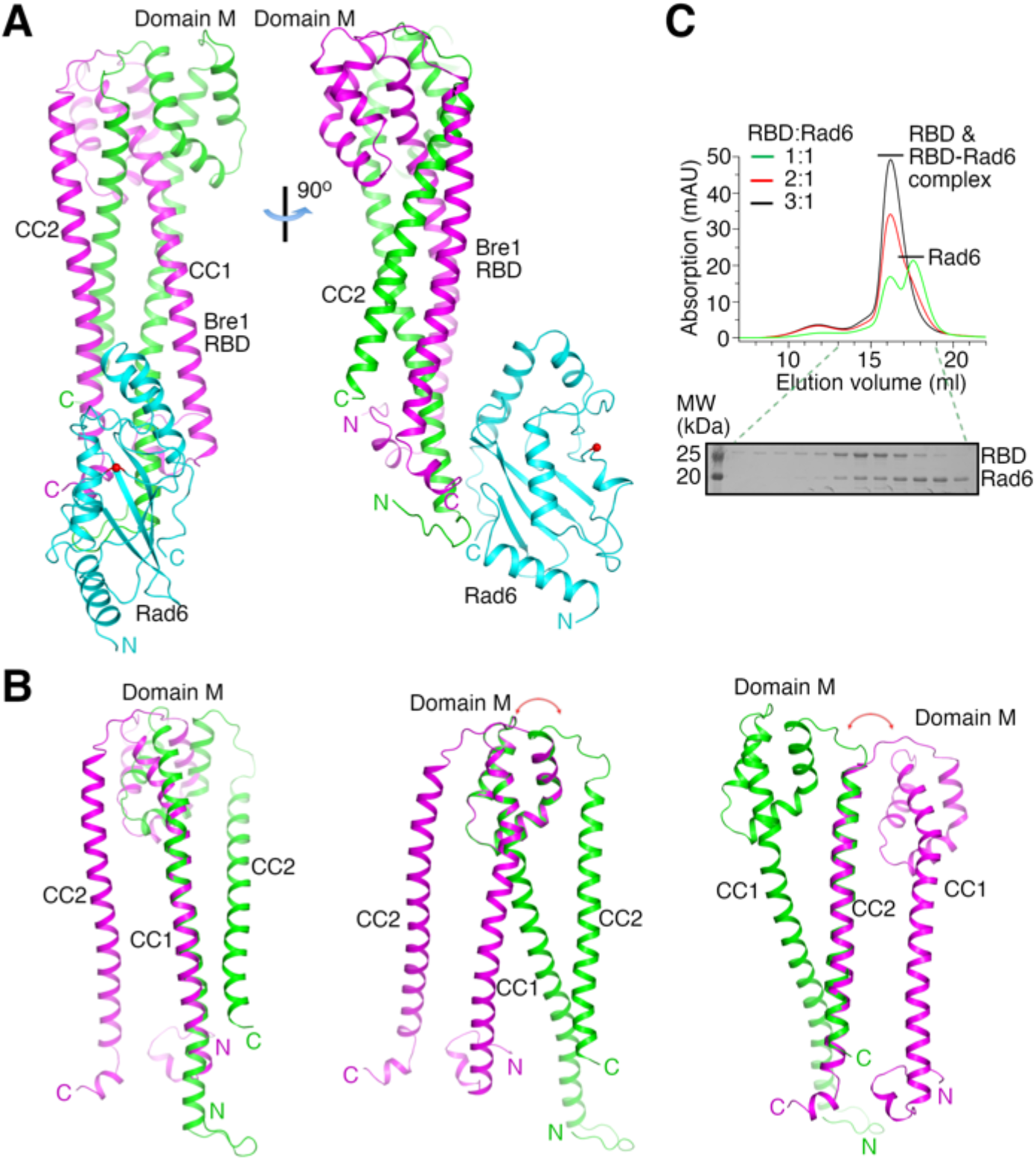
Bre1 RBD forms a 2:1 complex with Rad6. (A) Structure of the KlBre1 RBD-Rad6 complex. The two polypeptides in the KlBre1 RBD are colored in green and magenta, respectively. KlRad6 is colored in cyan. The red spheres indicate the KlRad6 active site. The N- and C-terminus of KlBre1 RBD and KlRad6 are indicated. This coloring scheme is used throughout the manuscript unless indicated otherwise. Structural figures are prepared with PyMOL (https://pymol.org/). (B) Structural comparison of the two polypeptides in the KlBre1 RBD. The CC1 (left), domain M (middle) or CC2 (right) regions in these polypeptides are aligned. The red arrows indicate the drastically different orientation of the domain M-CC2 linker in these polypeptides. (C) Gel filtration analysis of KlBre1 RBD and KlRad6 mixed at different molar ratios. 15 μM of KlRad6 was mixed with KlBre1 RBD with the indicated molar ratio, injected to a Superdex 200 10/300 column (GE healthcare) and eluted with a buffer containing 20 mM Tris (pH 7.5) and 200 mM sodium chloride. The lower panel shows SDS PAGE analysis of the gel filtration experiment with a 1:1 molar ratio. Source data is provided in figure 1 source data 1.

Both crystal forms 1 and 2 contain two KlBre1 RBD-Rad6 complexes in the asymmetric unit. The structures of KlBre1 RBD and KlRad6 in these four complexes are similar (Fig. S2A-B), except for regions surrounding KlRad6’s active site (detailed below). The structures of KlRad6 in both crystal forms are also similar to the previously reported structure of ScRad6(Worthylake *et al*., 1998) (Fig. S2B). The complexes in both crystal forms adopt similar conformations, except for complex 1 in crystal form 2. The orientation of KlRad6 in this complex is related to the orientation of KlRad6 in other complexes by a rotation of 14° (Fig. S2A). KlRad6 in this complex forms extensive interactions with neighboring protein molecules in the crystal (Fig. S2C), which may play a role in stabilizing the observed structure. In contrast, KlRad6 molecules in the other three complexes do not mediate extensive crystal packing interactions, their active sites are mostly exposed to solvent channels in the crystal. Their structure probably better reflect the structure of the KlBre1 RBD-Rad6 complex in solution. We will discuss this structure in the rest of the manuscript.

### Gel filtration characterization of the KlBre1 RBD-Rad6 complex

The dimeric structure of KlBre1 BRD is in line with a previous report(Kim & Roeder, 2009). To verify that KlBre1 RBD is dimeric in solution and forms a 2:1 complex with KlRad6, we performed gel filtration experiments. We found that the predominant species of KlBre1 RBD elutes at ~16 ml on a Superdex 200 10/300 column, almost the same as the KlBre1 RBD-Rad6 complexes used for crystallization (Figs S3A-B). Calibrating the column with molecules of known sizes indicated an apparent molecular weight of 80 kDa for KlBre1 RBD (Fig. S3C), somewhat larger than the expected molecular weight of the KlBre1 RBD dimer (52 kDa). The elongated shape of the KlBre1 RBD dimer may hinder its interaction with pores in the gel filtration medium, resulting in its earlier-than-expected elution. The KlBre1 RBD sample also contains some minor species that elute earlier. Sodium dodecyl sulphate-polyacrylamide gel electrophoresis (SDS PAGE) analysis did not reveal any major contaminating proteins (Fig. S3A), suggesting that they represent KlBre1 RBD species that fold differently or contain some minor contaminating proteins. KlRad6 elutes at ~17.5 ml on the same column, consistent with a monomeric form (Fig. S3D). When KlBre1 RBD and KlRad6 were mixed with a 1:1 molar ratio and applied to the column, a strong peak of free KlRad6 was observed (Fig. 1C), indicating that a significant amount of KlRad6 was in the free form. In contrast, when they were mixed with 2:1 or 3:1 molar ratio, this peak is absent (Fig. 1C). Together, these gel filtration experiments are consistent with a dimeric form of KlBre1 RBD in solution and a 2:1 KlBre1 RBD-Rad6 complex.

### Structural basis of the Bre1 RBD-Rad6 interaction

The structure indicates that KlRad6 interacts with the N-terminal region of KlBre1 RBD polypeptide 1 and N- and C-terminal regions of KlBre1 RBD polypeptide 2, which come together to form a single Rad6 binding site located at one end of the RBD. It interacts with the back side of KlRad6 opposite from its active site, composed of strands β1-β3, the C-terminus of α4 and neighboring loops (Figs 1A and 2A-B). Part of the C-terminal tail disordered in the structure of the free ScRad6(Worthylake *et al*., 1998) is structured in KlRad6 and mediate interactions with KlBre1 RBD (Figs 2B and S2B). Residues at the KlBre1 RBD-KlRad6 interface are mostly well-defined in the electron density map (Figs. S4A-B). The interface buries 2000 Å^2^ of surface area and contains hydrophobic, salt bridge and hydrogen bonding interactions (Figs 2A-B). At the interface, a hydrophobic patch consisting of KlBre1 residues Pro19, Leu20, Val25, Phe28, Val180’ (the prime sign indicates polypeptide 2 in KlBre1 RBD) and Phe181’ interacts with KlRad6 residues Ser20-Ser25, Met39, Pro43, Met153 and Met156; a second hydrophobic patch consisting of KlBre1 residues Ala32, Leu33, Cys36, Phe34’ and Leu37’ interacts with the KlRad6 Trp149 side chain; salt bridges are formed between Arg35 and Arg171’ in KlBre1 and Asp50 in KlRad6, Arg41’ in KlBre1 and Glu146 in KlRad6, and between Lys30’ in KlBre1 and Asp152 in KlRad6; hydrogen bonds are formed between the KlBre1 Gln29 side chain and the KlRad6 Thr52 side chain, the KlBre1 Gln29 and Lys30’ side chains and the KlRad6 Trp149 main chain carbonyl, and the KlBre1 Gln22 side chain and the KlRad6 Asp152 mainchain carbonyl.

**Figure 2.**
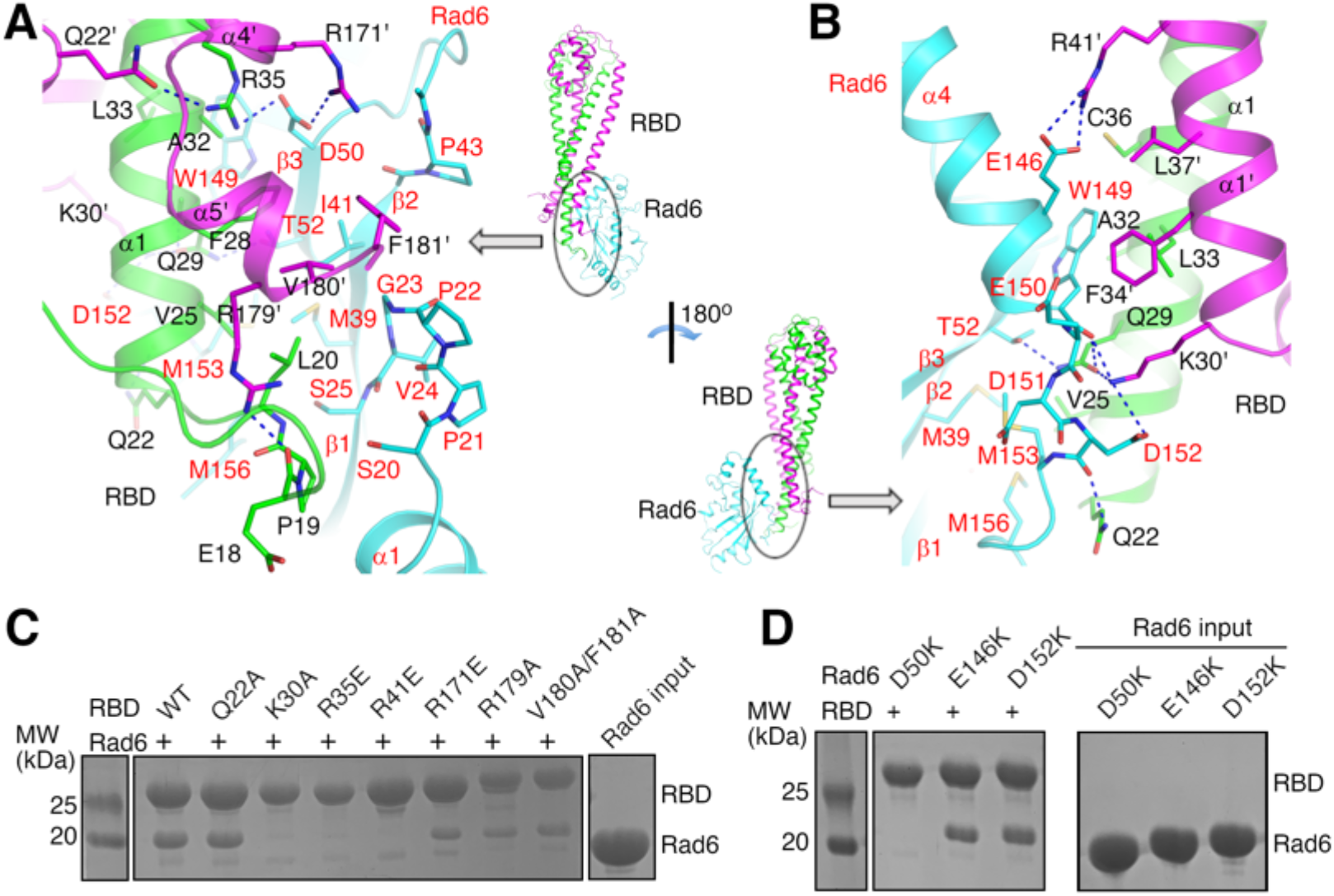
Structural basis of the KlBre1 RBD-Rad6 interaction. (A) and (B) Structure of the KlBre1 RBD-Rad6 interface. Residues important for the interaction are highlighted. The secondary structure elements for KlBre1 RBD are named as in Fig. S1. Labels for KlRad6 are in red. Labels with the prime sign are for the KlBre1 RBD polypeptide 2. (C) and (D) KlBre1 RBD-Rad6 pull down experiments. SDS PAGE analysis of strep-tagged KlBre1 RBD precipitated with strep-tactin beads and co-precipitated KlRad6 is shown. RBD, KlBre1 RBD. Source data for panels C and D are provided in figure 2 source data 1.

Several of the KlBre1 residues at the interface with KlRad6 also contribute to the interactions between the two KlBre1 polypeptides. These residues include Leu20, Leu33, Cys36, Phe34’, Leu37’ and Val180’, which mediate hydrophobic interactions between the two KlBre1 RBD polypeptides; and Gln29, Arg35 and Lys30’, which mediate hydrogen bond interactions (Figs 2A-B). In addition, several KlBre1 residues contributing to the interaction with KlRad6 in one polypeptide mediate the KlBre1 dimer interactions in the other polypeptide. These residues include Gln22’ (Fig. 2A) and Arg41 (Fig. S4C), which form hydrogen bonds with the Arg35 side chain and the Pro19’ mainchain carbonyl, respectively; and Phe34, Leu37, Pro19’, Leu33’ and Cys36’, which mediate hydrophobic interactions between the two KlBre1 RBD polypeptides (Fig. S4C).

To probe the function of important residues at the KlBre1 RBD-Rad6 interface, we introduced substitutions at these residues. In KlBre1 RBD, we introduced charge reversal substitutions R35E, R41E and R171E to disrupt the observed salt bridge interactions, and alanine substitutions at Gln22, Lys30, Val180 and Phe181 to disrupt the observed hydrogen bonding or hydrophobic interactions. We also introduced an R179A substitution to abolish the Arg179-Glu18 hydrogen bond, which stabilizes the Pro19-Leu20-containing loop for interaction with KlRad6 (Fig. 2A). In KlRad6, we introduced the D50K, E146K and D152K substitutions, which eliminate one of the salt bridges at the interface. Since some of the substitutions in KlBre1 RBD are located at the interface between the two KlBre1 polypeptides, we performed gel filtration experiments to test if they affect the dimerization of KlBre1 RBD. These experiments indicate that the substituted KlBre1 RBD eluted at the same volume as the wild type protein (Fig. S5A), suggesting that the substitutions do not alter the dimeric form. Similar gel filtration experiments indicated that the substitutions in KlRad6 do not significantly alter its overall structure (Fig. S5B).

We next probed the interaction with a pull-down experiment that precipitates streptavidin-binding peptide (strep-) tagged KlBre1 RBD with strep-tactin beads. A strong co-precipitation of KlRad6 was observed, consistent with its strong interaction with KlBre1 RBD (Fig. 2C). The co-precipitation was abolished by the K30A, R35E and R41E substitutions in KlBre1 RBD or the D50K substitution in KlRad6, but still present in experiments with the Q22A-, R171E-, R179A- or V180A/F181A-substituted KlBre1 RBD and the wild type KlRad6, or the wild type KlBre1 RBD and the E146K- or D152K-substituted KlRad6 (Figs 2C-D). To quantify the interaction and the effects of the substitutions, we performed isothermal titration calorimetry (ITC) experiments. ITC experiments performed with the same salt concentration as the pull-down experiments (200 mM) indicated a strong biphasic binding process (Fig. S6A). However, the heat exchange contributed by the minor phase is too small to enable reliable data fitting with a biphasic binding model but significant enough to interfere with data fitting with a monophasic binding model. We found that increasing the salt concentration to 1M suppressed the heat exchange contributed by the minor phase and enabled data fitting with a monophasic binding model, which gave a *K*_D_ of 12 nM (Fig. S6B and Table 2). The data fitting was not ideal, as the stoichiometry N-number is not close to the expected value of 0.5 for a 2:1 binding (Table 2). Nevertheless, since the data for the ITC experiment with 1M salt can be fitted, we proceeded to probe the effects of the substitutions on the KlBre1-KlRad6 interaction with this experiment. Consistent with the pull-down experiments, ITC experiments with 1M salt indicated that the K30A, R35E and R41E substitutions in KlBre1 RBD and the D50K substitution in KlRad6 reduced the affinity to undetectable levels (Figs S6B-C and Table 2). For substitutions that did not show strong effects in the pull-down experiments, including the Q22A, R179A and V180A/F181A substitutions in KlBre1 and the E146K and D152K substitutions in KlRad6, the ITC experiments indicated that they caused 17-640 folds increase in *K*_D_. Although the *K*_D_ values may not be accurate due to the non-ideal data fitting, these large increases in *K*_D_ indicate that the substitutions inhibited the KlBre1 RBD-Rad6 interaction to various degrees. The interaction between the R171E-substituted KlBre1 RBD and KlRad6 is undetectable in ITC experiments with 1M salt (Fig. S6B and Table 2), but quite substantial in pull-down experiments (Fig. 2C). We repeated the ITC experiment with 200 mM salt and found that the interaction was indeed detectable at this salt concentration (Fig. S6A and Table 2). The R171E substitution appears to have a stronger inhibitory effect on the KlBre1 RBD-Rad6 interaction at higher salt concentration. The pull-down and ITC experiments indicate that the substitutions we introduced in KlBre1 RBD and KlRad6 inhibit the interaction between them to various degrees. Together with the structure, these experiments provide a clear picture of the KlBre1 RBD-Rad6 interaction.

**Table 2.** Summary of ITC experiments

| KlBre1 RBD | KlRad6 | N | $K_D$ (μM) | $\Delta H$<br>(kcal/mol) | $T\Delta S$<br>(kcal/mol) | $\Delta G$<br>(kcal/mol) |
| --- | --- | --- | --- | --- | --- | --- |
| With 200 mM salt |  |  |  |  |  |  |
| WT | WT | * |  |  |  |  |
| R171E | WT | $0.047 \pm 0.366$ | $13.9 \pm 31.2$ | $-80 \pm 704$ | -73.4 | -6.63 |
| With 1M salt |  |  |  |  |  |  |
| WT | WT | $0.191 \pm 0.003$ | $0.012 \pm 0.009$ | $-30.0 \pm 1.32$ | -19.1 | -10.8 |
| Q22A | WT | $0.280 \pm 0.02$ | $0.207 \pm 0.182$ | $-18.3 \pm 2.38$ | -9.16 | -9.12 |
| K30A | WT | - | - | - | - | - |
| R35E | WT | - | - | - | - | - |
| R41E | WT | - | - | - | - | - |
| R171E | WT | - | - | - | - | - |
| R179A | WT | $0.142 \pm 0.007$ | $0.819 \pm 0.312$ | $-22.6 \pm 2.23$ | -14.3 | -8.31 |
| V180A/F181A | WT | $0.193 \pm 0.005$ | $7.64 \pm 0.903$ | $-31.6 \pm 1.78$ | -24.6 | -6.98 |
| WT | D50K | - | - | - | - | - |
| WT | S111L | $0.289 \pm 0.004$ | $0.007 \pm 0.008$ | $-33.6 \pm 1.52$ | -22.5 | -11.1 |
| WT | E146K | $0.134 \pm 0.002$ | $0.138 \pm 0.021$ | $-32.3 \pm 0.63$ | -22.9 | -9.37 |
| WT | D152K | $0.134 \pm 0.002$ | $0.827 \pm 0.105$ | $-30.4 \pm 1.02$ | -22.1 | -8.30 |

\* Unreliable data fitting

Previous mutagenesis studies indicated that substitutions on Lys31 in ScBre1 and on Gly23 and Asp50 in ScRad6 abolished their binding(Turco *et al*., 2015). In our structure the equivalent residues, Lys30 in the KlBre1 RBD and Gly23 and Asp50 in KlRad6, mediate important interactions between KlBre1 RBD and KlRad6 (Figs 2A-B). Therefore, the reported loss of binding is most likely due to the loss of important interactions at the ScBre1 RBD-Rad6 interface.

### The Bre1 RBD-Rad6 interaction stimulates free ubiquitin chain production by Rad6

Ubiquitin has been reported interact with the back side of several E2 enzymes(Brzovic *et al*, 2006; Buetow *et al*, 2015; Eddins *et al*, 2006; Hibbert *et al*, 2011; Sakata *et al*, 2010). The activity of some of these E2 enzymes, including the human Rad6B, is stimulated by such interaction(Brzovic *et al*., 2006; Buetow *et al*., 2015; Hibbert *et al*., 2011). The expected ubiquitin binding site on Rad6’s back side overlaps with the Bre1 RBD binding site (Fig. 3A, left panel), promoting us to probe the effect of Bre1 RBD on Rad6’s activity. Rad6 possesses an intrinsic activity to catalyze free ubiquitin chain formation(Hibbert *et al*., 2011). We probed the effect of Bre1 RBD on this activity. To assess the basal activity of Rad6, we introduced a G23R/T52A substitution to KlRad6’s back side, the equivalent of which has been reported to abolish ubiquitin binding to the back side of the human Rad6B and the stimulatory effect it mediates(Hibbert *et al*., 2011). Gel filtration experiments suggest that the G23R/T52A substitution did not alter the overall structure of KlRad6 (Fig. S5B). Consistent with ubiquitin binding to the back side of KlRad6 and stimulating its activity, we found that the G23R/T52A substitution significantly hindered the free ubiquitin chain production by KlRad6 (Figs 3B and S7A). Supplementing KlBre1 RBD to the reaction with the wild type KlRad6 significantly accelerated the formation of ubiquitin chains (Fig. S7B), but the ubiquitin chains can be attached to KlBre1 RBD (Fig. S7C). To access the amount of free ubiquitin chains, we removed KlBre1 RBD and ubiquitin chains attached to it after the reaction with strep-tactin beads, which binds to the strep-tagged KlBre1 RBD. After this treatment, we found that the reaction with the wild type KlRad6 and KlBre1 RBD also produced significantly more free ubiquitin chains than the reaction with the G23R/T52A-substituted KlRad6 (Fig. 3B). The affinity between ubiquitin and Rad6 is rather weak, with *K*_D_ in the mM range(Kumar *et al*, 2015). In contrast, our binding experiments indicated a strong interaction between KlBre1 RBD and KlRad6. Therefore, when KlBre1 RBD is supplemented to the reaction, KlBre1 RBD but not ubiquitin is expected to occupy KlRad6’s back side. Thus, our experiments indicate that KlBre1 RBD stimulates the free ubiquitin chain production by KlRad6.

**Figure 3.**
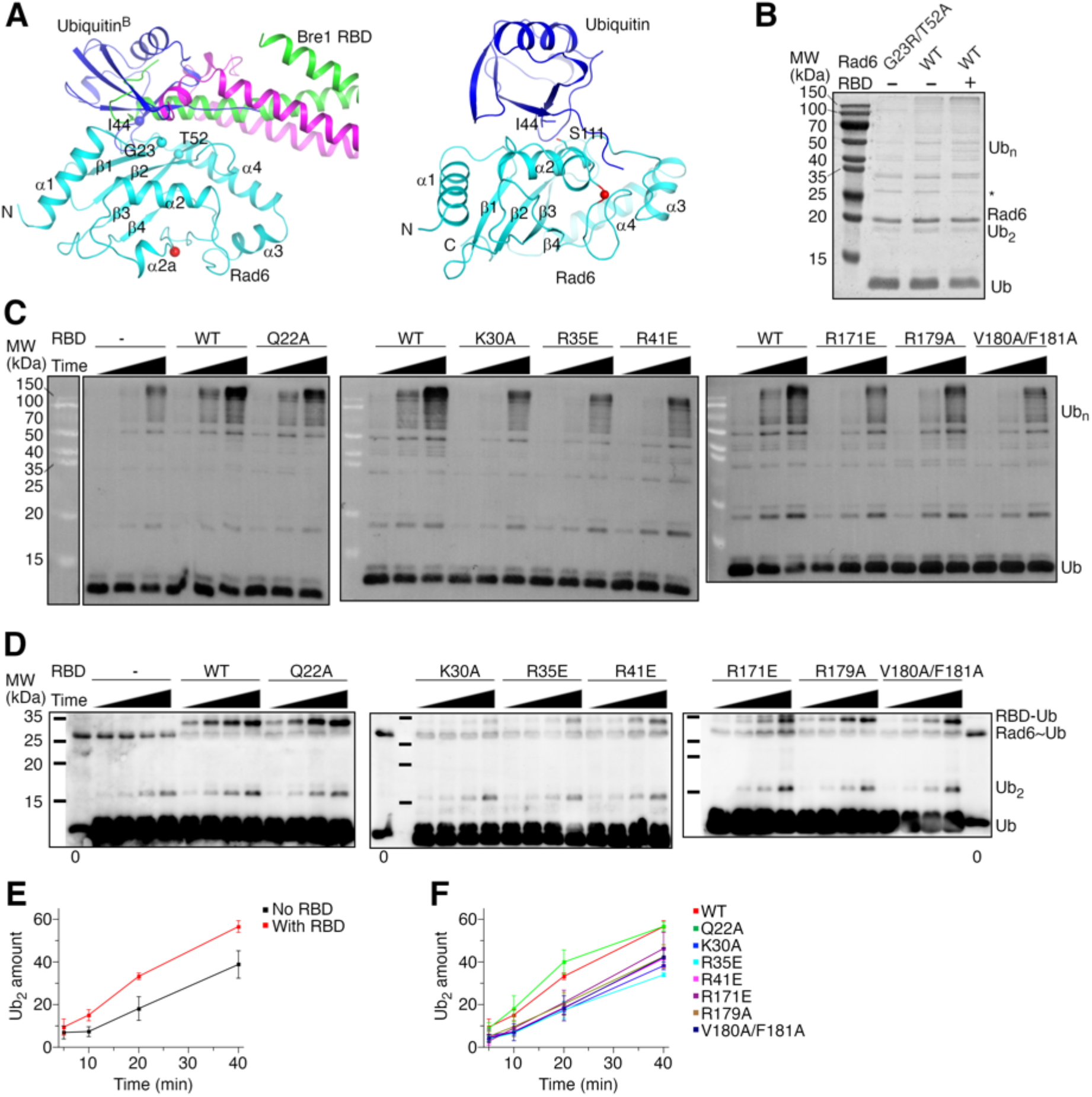
KlBre1 RBD stimulates KlRad6’s enzymatic activity. (A) Structural modeling of the ubiquitin-KlRad6 interaction. The left panel shows a model of KlRad6 with ubiquitin bound at its back side (ubiquitin^B^). The model is based on the structure of UbcH5c with ubiquitin bound at its back side (PDB 2FUH)(Brzovic *et al*., 2006). KlBre1 RBD observed in our structure is shown for reference. The right panel shows a model of the KlRad6~ubiquitin conjugate in the closed conformation. The model is based on the closed conformation of the ubiqtuin-Ubc13 covalent complex (PDB 5ait)(Branigan *et al*, 2015). Secondary structural elements and the N- and C-termini of KlRad6 are indicated. Important residues are highlighted. The red spheres indicate KlRad6’s active site. (B) SDS PAGE analysis of free ubiquitin chain production by KlRad6. The reactions were carried out for 30 minutes. KlBre1 RBD and ubiquitin chains attached to it were removed before analysis. The star sign indicates the position of KlBre1 RBD, which is mostly removed (compare to Fig. S7B). Ub, ubiquitin; Ub_2_ and Ub_n_, ubiquitin chains with 2 or n ubiquitin moieties, respectively. (C) Western blot analysis of free ubiquitin chain production in reactions with I44A-substituted ubiquitin and S111L-substituted KlRad6. Reactions carried out for 5, 10 and 20 minutes are presented. KlBre1 RBD and ubiquitin chains attached to it were removed before analysis. Two independent repeats of the experiments were performed. (D) Western blot analysis of the Rad6~ubiquitin discharging reaction. K0/I44A-substituted ubiquitin charged to S111L-substituted KlRad6 was discharged to I44A-substituted ubiquitin in the absence or presence of KlBre1 RBD. The reactions were allowed to proceed for 5, 10, 20 and 40 minutes and analyzed by non-reducing SDS PAGE followed by western blot for ubiquitin. In lanes marked with “0”, western blot analysis of the KlRad6~ubiqtuin conjugate prior to the discharging reaction is presented. The band positions of KlBre1 RBD and the KlRad6~ubiquitin conjugate are almost identical in SDS PAGE (Fig. S7F). KlBre1 RBD may interfere with anti-body binding to the KlRad6~ubiqutin conjugate, resulting in its poor detection in western blot. Three independent repeats of the experiments were performed. (E) and (F) Quantification of the di-ubiquitin production in KlRad6~ubiquitin discharging reactions. The intensity of the di-ubiquitin band divided by the intensity of the Rad6~ubiqtuin band in lane “0” of the same blot times 100 was calculated to represent the di-ubiquitin amount. The error bars represent standard deviations of three independent experiments. Band intensities were read with ImageJ. Source data for panels B-D are provided in figure 3 source data 1, for panels E and F are provided in figure 3 source data 2.

Consistent with the role of KlRad6 residues Gly23 and Thr52 in mediating interactions with KlBre1 RBD (Figs 2A-B), we found a severe inhibition of the KlBre1 RBD-Rad6 interaction by the G23R/T52A substitution (Fig. S7D). Hence, the G23R/T52A-substituted KlRad6 is not expected to be activated by KlBre1 RBD, preventing a comparison of the basal and Bre1 RBD-stimulated activities of the same Rad6 variant. To do so, we utilized the I44A-substituted ubiquitin to assess the basal activity of KlRad6. Ile44 in ubiquitin mediate important interactions with the back side of E2 enzymes (Fig. 3A, left panel) and the interaction is inhibited by the I44A substitution(Brzovic *et al*., 2006; Buetow *et al*., 2015; Eddins *et al*., 2006; Hibbert *et al*., 2011; Sakata *et al*., 2010). The substitution also destabilizes the closed conformation of the E2~ubiquitin conjugate required for ubiquitin transfer, but this defect can be rescued by substitutions in the E2 enzyme near the active site(Li *et al*, 2015; Saha *et al*, 2011). One of such substitutions, equivalent to S111L in KlRad6, introduces hydrophobic interactions with the substituted I44A side chain to stabilize the closed conformation (Fig. 3A, right panel). The S111L substitution did not inhibit the KlBre1 RBD-Rad6 interaction (Figs S6C, S7D and Table 2) or alter the overall structure of KlRad6 (Fig. S5B). Consistent with it stabilizing the closed conformation of the KlRad6~ubiquitin (I44A) conjugate, it rescued the severely inhibited free ubiquitin chain production associated with the I44A-substituted ubiquitin (Fig. S7A). Supplementing KlBre1 RBD into the reaction with the S111L-substituted KlRad6 and I44A-subsituted ubiquitin significantly stimulated the free ubiquitin chain production, consistent with a stimulatory effect of KlBre1 RBD on free ubiquitin chain production by KlRad6 (Fig. 3C).

To further probe the role of the KlBre1 RBD-Rad6 interaction in the stimulation of KlRad6’s activity, we tested the effects of KlBre1 substitutions that inhibit the interaction. In free ubiquitin chain formation reactions with S111L-substituted KlRad6 and I44A-subsituted ubiquitin, we found that the Q22A substitution did not noticeably change the stimulation by KlBre1 RBD, and substitutions K30A, R35E, R41E, R171E, R179A and V180A/F181A significantly inhibited stimulation (Fig. 3C). These observations correlate well with our binding experiments, which indicate that the Q22A substitution moderately inhibited the KlBre1 RBD-KlRad6 interaction and caused a 17-fold increase in *K*_D_, whereas the other substitutions caused much stronger inhibitions (Figs 2C, S6B and Table 2). Together, these data indicate a critical role of the Bre1 RBD-Rad6 interaction in stimulating Rad6’s activity.

### The Bre1 RBD-Rad6 interaction stimulates Rad6’s enzymatic activity in ubiquitin discharging

The ubiquitination reaction consists of two steps, including ubiquitin charging to the E2 enzyme to form the E2~ubiquitin conjugate, and ubiquitin discharging from the conjugate to the substrate. Ubiquitin and several E3 enzymes have been reported to bind to the back side of E2 enzymes to regulate the discharging reaction(Buetow *et al*., 2015; Das *et al*, 2009; Li *et al*., 2015; Metzger *et al*, 2013). To test if the Bre1 RBD-Rad6 interaction affects ubiquitin discharging from the Rad6~ubiquitin conjugate, we performed single turnover ubiquitin discharging experiments. After charging ubiquitin to Rad6 with the E1 enzyme and ATP, we stopped the charging reaction and allowed the KlRad6~ubiquitin conjugate to dis-integrate. We found that the amount of the conjugate decreases over time and adding KlBre1 RBD significantly accelerated the rate of decrease (Fig. S7E). Since KlBre1 RBD itself can be ubiquitinated, it could accelerate the discharging reaction by increasing the amount of substrate. To eliminate this factor and test if KlBre1 RBD regulates the enzymatic activity of KlRad6 in ubiquitin discharging, we monitored the reaction of ubiquitin discharging to ubiquitin, the substrate amount of which is unaffected upon supplementing KlBre1 RBD. We monitored the amount of the reaction product, the free ubiquitin chain. To enhance the product signal, we substituted all lysine residues in the donor ubiquitin to arginine (K0 substitution). After its attachment to the acceptor ubiquitin, additional ubiquitin attachment to the K0-substituted ubiquitin is prevented, making di-ubiquitin the only possible product. To eliminate the potential regulatory effect of ubiquitin on KlRad6 by binding to its back side, we introduced the I44A substitution to ubiquitin and the S111L substitution to KlRad6. We found that the di-ubiquitin production in this reaction is significantly stimulated by the wild type KlBre1 RBD (Figs 3D-E). In addition, we found that the Q22A substitution in KlBre1 RBD that moderately inhibits its interaction with Rad6 had little effect on the stimulation, whereas substitutions K30A, R35E, R41E, R171E, R179A and V180A/F181A that strongly inhibited the interaction strongly inhibited the stimulation (Figs 3D and F). Together, these data indicate that the KlBre1 RBD-Rad6 interaction stimulates KlRad6’s enzymatic activity in ubiquitin discharging from the KlRad6~ubiquitin conjugate.

### Bre1 RBD increases Rad6’s active site accessibility

Analysis of our structure provided insights into the mechanism of the observed Rad6 stimulation. When the three KlRad6 molecules not mediating extensive crystal packing interactions in our structures are compared, large structural differences are observed residues 90-98 and 115-121 surrounding KlRad6’s active site (Fig. 4A). The 90-98 region can be resolved in the electron density map in crystal form 1 but is mostly disordered in KlRad6 molecule 2 in crystal form 2. The structures of the 115-121 region in these molecules are significantly different, the largest distance between the equivalent Cα atoms in this region is 5.2 Å. These regions in the three KlRad6 molecules also display significantly elevated temperature factors (Fig. 4B). The average temperature factor for these regions in KlRad6 molecules 1 and 2 in crystal form 1 and molecule 2 in crystal form 2 are 143.7 Å^2^, 156.4 Å^2^ and 173.5 Å^2^, respectively; whereas the average temperature factor for the rest of these molecules are 104.6 Å^2^, 117.1 Å^2^ and 149.2 Å^2^. The temperature factor of these regions in KlRad6 molecule 1 in crystal form 2 is not elevated (Fig. S8A), consistent with their role in mediating extensive crystal packing interactions (Fig. S2C). The equivalent regions in Rad6 orthologs have been reported to be mobile. NMR studies indicated that these regions in the human Rad6B are flexible on a nanosecond to picosecond time scale(Miura *et al*, 1999; Miura *et al*, 2002), structural differences were also observed in these regions in the crystal structure of the closely related ScRad6(Worthylake *et al*., 1998), which shares 94.5% sequence identity with KlRad6. However, the structural difference in the crystal structure of ScRad6 is not as pronounced. Three ScRad6 molecules are present in the crystal, their 90-98 region adopt almost identical structures, the largest distance between the equivalent Cα atoms in the 115-121 region is 4.1 Å (Fig. 4C). The temperature factor of these regions is also elevated in this structure, but much less significantly (Figs 4D and S8B). The average temperature factor for these regions in the three ScRad6 molecules are 45.7 Å^2^, 39.8 Å^2^ and 44.5 Å^2^, and the average temperature factor for the rest of these molecules are 42.6 Å^2^, 34.7 Å^2^ and 38.1 Å^2^. Together, these structural data indicate an increased mobility of regions 90-98 and 115-121 in KlRad6 in the KlBre1 RBD-Rad6 complex, suggesting that the KlBre1 RBD allosterically increase the mobility of these regions. Such increase in mobility is expected to make KlRad6’s active site more accessible, which facilitates the nucleophilic attack of the KlRad6~ubiqutin thioester bond and promotes ubiquitin discharge from the KlRad6~ubiquitin conjugate.

**Figure 4.**
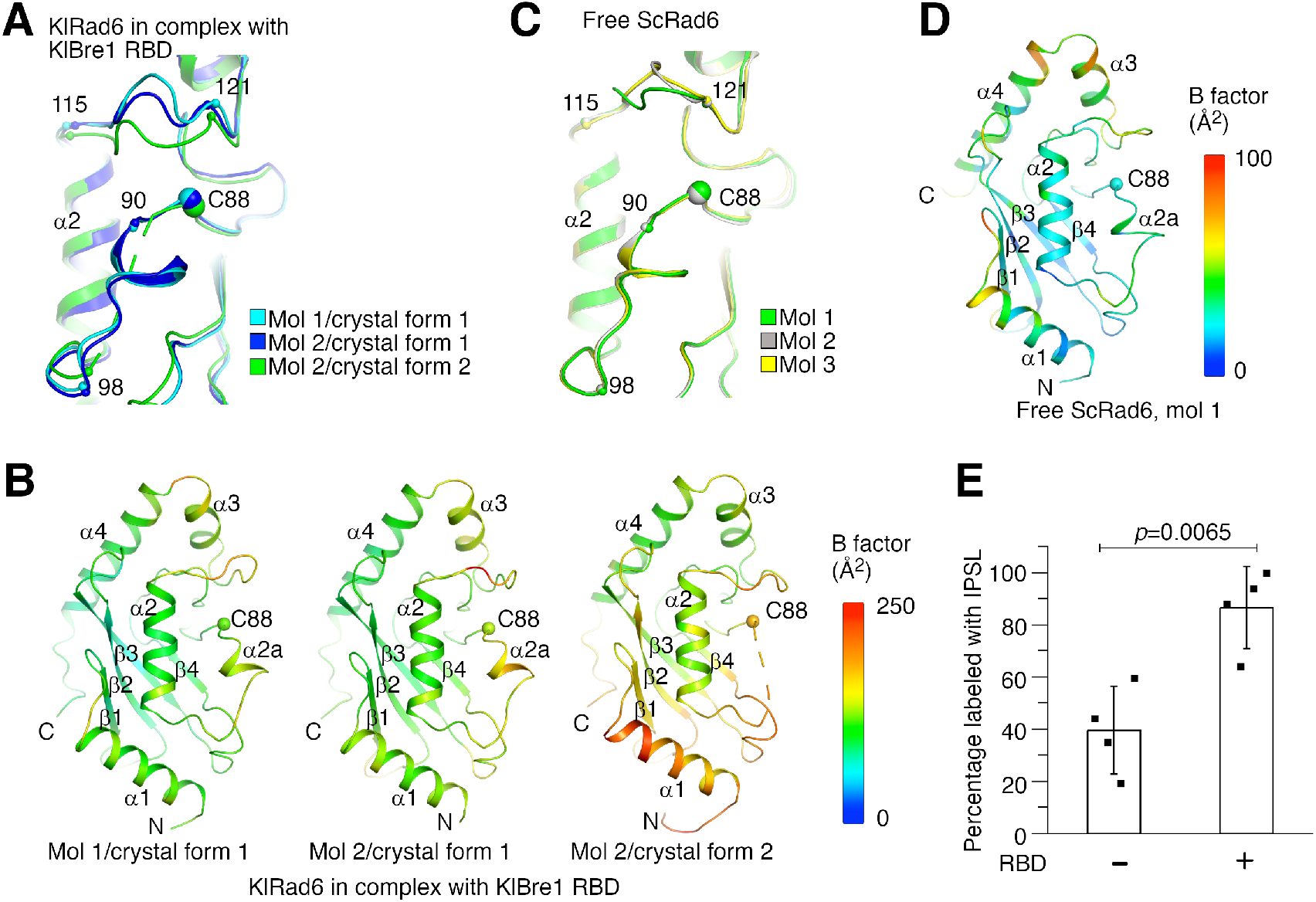
Bre1 RBD increases Rad6’s active site accessibility. (A) Structural comparison of KlRad6 molecules in our crystals. Regions surrounding the active site (C88) in KlRad6 molecules not mediating extensive crystal packing interactions are shown. The Ca positions of residues 90, 98, 115 and 121 are indicated. (B) Temperature factor distribution of KlRad6 molecules in our crystals. The structures of KlRad6 molecules not mediating extensive crystal packing interactions are colored according to the temperature factor. (C) Structural comparison of ScRad6 molecules in the crystal structure of free ScRad6 (PDB 1AYZ)(Worthylake *et al*., 1998). (D) Temperature factor distribution of ScRad6 in the crystal structure of free ScRad6. Molecule 1 in the crystal is presented. (E) KlBre1 RBD increase KlRad6’s active site accessibility. IPSL labeling of KlRad6’s active site cysteine is presented. The error bars represent standard deviations of four independent experiments (represented by the black dots). The *p* value is derived from the two-sided Student’s *t*-test. Source data is provided in figure 4 source data 1.

To test the above possibility, we probed the accessibility of KlRad6’s active site cysteine with 3-(2-Iodoacetamido)-2,2,5,5-tetramethyl-1-pyrolidinyloxy (IPSL), a cysteine-reacting reagent. We found that after a 15-minute incubation with 300 μM IPSL, 39% of KlRad6’s active site cysteine was labeled with IPSL, supplementing KlBre1 RBD drastically increased the labeling level to 86% (Fig. 4E). The drastic increase in IPSL labeling together with the structural data indicate that KlBre1 RBD allosterically increases KlRad6’s active site accessibility to stimulate its activity.

### The Bre1 RBD-Rad6 interaction is crucial for Bre1’s function inside the cell

To assess the physiological function of the Bre1 RBD-Rad6 interaction inside the cell, we generated a *BRE1* knock out yeast strain and complemented it with the wild type or substituted ScBre1. The effects of several ScBre1 substitutions were assessed, including Q23A, Q30A/K31D and R36E/R42E. In line with previous reports(Hwang *et al*., 2003; Wood *et al*., 2003), we found that knocking out *BRE1* abolished H2Bub1 *in vivo* in unperturbed cells and in cells under hydroxy urea (HU)-induced replication stressed conditions (Fig. 5A), and caused a strong sensitivity towards replication stress or DNA damaging reagents including HU, camptothecin (CPT), methylmethane sulfonate (MMS) and phleomycin (Fig. 5B). These phenotypes can be rescued by complementing the cells with the wild type ScBre1, or ScBre1 with the Q23A substitution, but not ScBre1 with the Q30A/K31D or R36E/R42E substitutions. The Bre1-mediated H2Bub1 plays a critical role in promoting DNA DSB repair by HR(Zheng *et al*., 2018). To test the importance of the Bre1 RBD-Rad6 interaction in HR, we employed an ectopic recombination system in which a single DSB is generated by the HO endonuclease at the *MAT**a*** sequence inserted in chromosome V, which can be repaired by HR using the homologous *MAT***a**-inc sequence located on chromosome III (Fig. 5C)(Ira *et al*, 2003). As previously reported, over 80% of the cells with the wild type *BRE1* completed the repair and survived, knocking out *BRE1* reduced the survival rate to less than 50% (Fig. 5D). Remarkably, the HR defect of *BRE1* knock out cells can be rescued by introducing a plasmid-borne *bre1-Q23A* allele but not the *bre1-Q30A/K31D* or *bre1-R36E/R42E* allele (Fig. 5D). These data are in line with our structure and biochemical studies. The Q23A substitution did not cause noticeable defects in the cellular H2Bub1 level, tolerance towards replication stress or DNA damaging reagents or the HR repair. The equivalent substitution in KlBre1 RBD, Q22A, only moderately reduced its affinity towards KlRad6. In contrast, strong defects were observed for the Q30A/K31D and R36E/R42E substitutions. KlBre1 residues equivalent to the substituted ones mediate important hydrogen bond or salt bridge interactions with Rad6 (Gln29, Lys30, Arg35 and Arg41, Figs 2A-B), and our binding experiments indicated that substitutions on these residues abolished the KlBre1 RBD-KlRad6 interaction (Figs 2C, S6B and Table 2). The strong defects associated with these substitutions correlates with the expected loss of the ScBre1 RBD-Rad6 interaction. Together, these data indicate that the Bre1 RBD-Rad6 interaction is crucial for the Bre1-mediated H2B mono-ubiquitination, DNA damage response and HR repair inside the cell.

**Figure 5.**
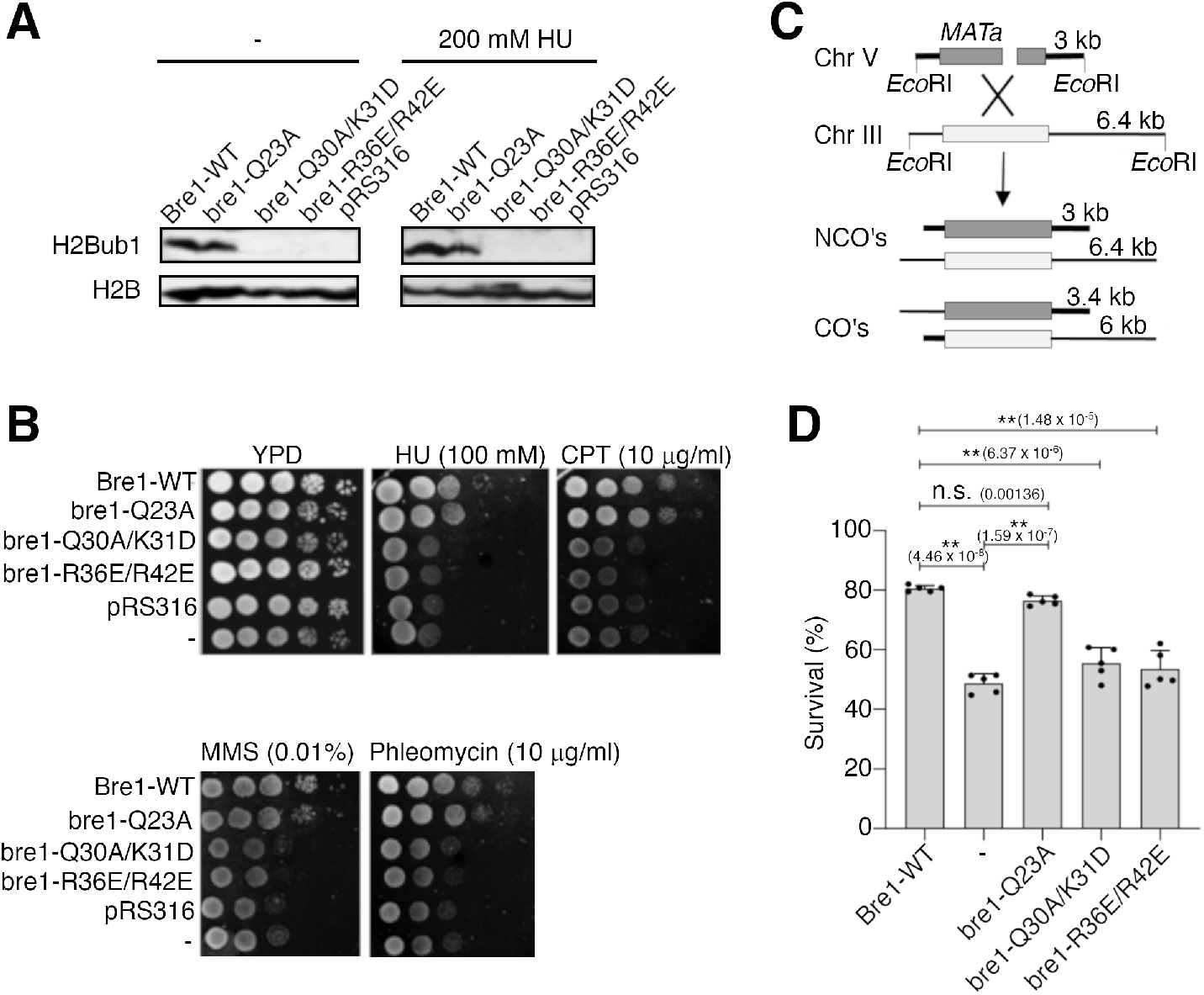
The Bre1 RBD-Rad6 interaction is crucial for Bre1’s function inside the cell. (A) The Bre1 RBD-Rad6 interaction is essential for the H2Bub1 formation inside the cell. Western blot analysis for H2Bub1 in *BRE1* knock out cells complemented with pRS316-derived plasmids carrying the wild type or substituted ScBre1 is shown. Data for cells complemented with the empty vector are included for comparison. Cells were unperturbed (-) or stressed with 200 mM HU. Source data is provided in figure 5 source data 1. (B) The Bre1 RBD-Rad6 interaction is essential for survival in the presence of replication stress/DNA damaging reagents. Growth of *BRE1* knock out cells complemented with pRS16-derived plasmids for the wild type or substituted ScBre1 is shown. The cells were challenged with HU, CPT, MMS or phleomycin. Data for cells complemented with the empty vector or uncomplemented cells (-) are included for comparison. (C) Scheme showing the ectopic recombination system. CO, crossover; NCO, non-crossover. (D) The Bre1 RBD-Rad6 interaction is essential for the HR repair. Survival rate of *BRE1* knock out cells complemented with the wild type or substituted ScBre1 due to successful HR repair is shown. Data for the wild type or uncomplemented cells (-) are included for comparison. Error bars represent standard deviations of five independent experiments (represented by the black dots). ** indicates *p*<0.001 in the two-sided Student’s *t*-test. n.s., not significant. The *p* values are indicated in the brackets. Source data is provided in figure 5 source data 2.

## Discussion

Our structure and binding studies provide mechanistic insights into the interaction between Bre1 RBD and Rad6. We found that Bre1 RBD forms a dimer and multiple regions in both polypeptides contribute to interaction with Rad6 to form a stable Bre1 RBD-Rad6 complex. Bre1 has been reported to be dimeric and its N-terminal region plays a critical role in its dimer formation(Kim & Roeder, 2009). Our data indicate that the Bre1 dimerization is required for the formation of a functional RBD.

An important finding we made is that the Bre1 RBD-Rad6 interaction stimulates Rad6’s enzymatic activity. Our data indicate that by binding to Rad6’s back side, Bre1 RBD allosterically increases the accessibility of Rad6’s activity site, facilitating the nucleophilic attack of the Rad6~ubiqtuin thioester bond to promote the ubiquitin discharging. Ubiquitin also binds to the back side of several E2 enzymes to stimulate their activity(Brzovic *et al*., 2006; Buetow *et al*., 2015; Hibbert *et al*., 2011). However, there are striking differences between the Bre1 RBD-mediated and ubiquitin-mediated stimulation. Unlike Bre1 RBD, ubiquitin does not form stable complexes with these E2 enzymes, and it is not clear if it allosterically increases their active site accessibility. In addition to the above mechanism, our data suggests that the Bre1 RBD-Rad6 interaction may promote the H2Bub1 catalysis through additional mechanisms. First, it may maintain a close Bre1-Rad6 association during the H2Bub1 catalysis. Structural studies indicated that the E1 enzyme and the RING domain in the RING family of E3 enzymes compete for binding to the E2 enzyme(Cappadocia & Lima, 2018; Gundogdu & Walden, 2019; Stewart *et al*., 2016; Streich & Lima, 2014), resulting a “ping-pong” motion of the E2 enzyme between the E1 enzyme and the RING domain during the ubiquitination catalysis. The Bre1 RBD-Rad6 interaction does not interfere with the expected Rad6-E1 or Rad6-RING interactions (Figs S9A-B). Therefore, the strong affinity between Bre1 RBD and Rad6 we and others(Turco *et al*., 2015) observed suggests that Rad6 remain associated with Bre1 during the ubiquitination catalysis by binding to its RBD. Such a Rad6-Bre1 association could increase the local concentration of Rad6 to promote the catalysis. Second, the Bre1 RBD-Rad6 interaction may also stabilize the closed conformation of the bound Rad6~ubiquitin conjugate for ubiquitin transfer. Structural modeling suggests that Bre1 RBD could interact with ubiquitin in the conjugate to stabilize its closed conformation, the side chain of the conserved Phe181 in KlBre1 is expected to play a central role (Fig. S9C). The loss of this interaction may contribute to the observed loss of KlRad6 stimulation by the V180A/F181A-substituted KlBre1 RBD (Figs 3C, D and F). These important functions of the Bre1 RBD-Rad6 interaction are in line with our observation that it is crucial for multiple H2Bub1-regulated processes inside the cell.

In a recent study, it was found that supplementing ScBre1 RBD to a single turnover discharging experiment stimulates ubiquitin discharging from the Rad6~ubiquitin conjugate(Turco *et al*., 2015). Since ScBre1 RBD can probably be ubiquitinated in such experiments like KlBre1 RBD, it could contribute to the observed stimulation by increasing the amount of substrate. Therefore, it is difficult to conclude from the reported data whether ScBre1 RBD directly stimulates Rad6’s enzymatic activity, which is relevant to the H2Bub1 catalysis *in vivo*. In our study, we monitored the reaction of ubiquitin discharging to ubiquitin, whose substrate amount is unaffected upon supplementing Bre1 RBD. We found that the reaction rate is significantly stimulated by supplementing KlBre1 RBD and the stimulation is inhibited by substitutions that inhibits the KlBre1 RBD-Rad6 interaction. Together with the observation that KlBre1 RBD stimulates free ubiquitin chain production by Rad6, our data provided strong evidence that the Bre1 RBD-Rad6 interaction stimulates Rad6’s enzymatic activity.

Several other E3 enzymes have been reported to contain specific E2 enzyme back side binding regions (E2BBRs), which regulate the activity of related E2 enzymes with distinct mechanisms. E2BBRs in Gp78(Das *et al*, 2013; Das *et al*., 2009) and Cue1(Metzger *et al*., 2013) have been reported to stimulate the activity of the related E2 enzymes by increasing their active site accessibility and interaction with the RING domain, whereas E2BBRs in AO7(Li *et al*., 2015) and Rad18(Hibbert *et al*., 2011) have been reported to inhibit the activity of the related E2 enzymes by blocking ubiquitin binding to their back side and the stimulatory effect it mediates. All the previously reported E2BBRs possess a single E2-interacting region that contains a major a helix, which bind to the back side of E2 enzymes in different orientations (Fig. S10). In sharp contrast to these E2BBRs, multiple regions in both polypeptides in the Bre1 RBD dimer contribute to the interaction with the Rad6 back side. Most interestingly, although both the Bre1 RBD and the Rad18 E2BBR bind to the Rad6 back side, they appear to have opposite effects on Rad6’s activity. Together, these previously studies and ours indicate that E3 enzymes can interact with the back side of E2 enzymes with drastically different mechanisms to regulate their activity.

In the Bre1 holoenzyme, RBD coordinates with other domains in Bre1 and additional factors to catalyze the highly specific and efficient H2Bub1 reaction. Our study provided glimpse into this important reaction, yet extensive future studies are required to fully understand its mechanism. One of the most important questions to answer is how Bre1 RBD coordinates with its RING domain during the H2Bub1 catalysis. Unlike Bre1 RBD, the Bre1 RING domain appears to be a symmetrical dimer with two Rad6 binding sites(Kumar & Wolberger, 2015), although probably only one of them interacts with the Rad6~ubiquitin conjugate for ubiquitin transfer to NCP(Gallego *et al*., 2016). It is also important to understand the coordination between Bre1 RBD and additional domains/factors during the H2Bub1 catalysis. For instance, it was recently reported that ubiquitin interacts with a non-canonical back side of Rad6 to regulate its activity(Kumar *et al*., 2015), which appears to be accessible in the Bre1 RBD-bound Rad6 (Fig. S9B). A clear understanding of the H2Bub1 catalytic mechanism will help to resolve the long-standing puzzle how Bre1 directs Rad6’s activity towards substrate mono-ubiquitination. Rad6 participates in both ubiquitin chain formation and mono-ubiquitination reactions. It functions with the E3 enzyme Ubr1 to catalyze the ubiquitin chain modification of N-end rule protein substrates(Dohmen *et al*, 1991), with Bre1(Hwang *et al*., 2003; Robzyk *et al*., 2000; Wood *et al*., 2003) and Rad18(Hoege *et al*, 2002) to mono-ubiquitinate H2B and the proliferating cell nuclear antigen (PCNA), respectively. It also possesses an intrinsic activity to catalyze free ubiquitin chain formation(Hibbert *et al*., 2011). The Rad18 E2BBR suppresses its intrinsic free ubiquitin chain forming activity and directs its activity towards PCNA mono-ubiquitination(Hibbert *et al*., 2011). In contrast, our data indicates that Bre1 RBD does not inhibit the free ubiquitin chain formation by Rad6 but stimulates its activity. Thus, Bre1 utilizes a completely different mechanism to direct Rad6’s activity towards H2B mono-ubiquitination. Understanding this mechanism is essential for a clear picture of the H2Bub1 catalysis and will provide insights into the catalytic mechanism of E3 enzymes in general.

The human orthologs of Bre1, RNF20 and RNF40, form a heterodimer(Kim *et al*., 2009). It has been reported that the RNF20 N-terminal 381 residues could form a complex with RNF40 that binds Rad6(Kim *et al*., 2009), suggesting that these residues participate in the formation of a functional RBD. However, sequence analysis suggested that the region spanning residues 328-528 in RNF20 possesses homology to the RBD region in Bre1(Zhu *et al*., 2005). Therefore, future studies are required to define the RBD in the RNF20/RNF40 heterodimer and elucidate its structure and function.

## Materials and Methods

### Protein expression and purification

The coding regions for the *Kluyveromyces lactis* Bre1 (KlBre1) fragments 1-206 or 1-184 and the full length Rad6 (KlRad6) were amplified from the *Kluyveromyces lactis* genome and inserted into vectors pET26B and pET28A (Novagen), respectively. The recombinant KlBre1 fragments and KlRad6 contain no tags and an N-terminal 6x histidine (his-) tag, respectively. For protein expression, *Escherichia coli* BL21 Rosetta (DE3) cells harboring these plasmids were induced with 0.2 mM isopropyl β-D-1-thiogalactopyranoside (Bio Basic) for 16 hours at 16 °C. For complex purification, cells expressing one of the KlBre1 fragments were mixed with cells expressing KlRad6 and lysed with an AH-2010 homogenizer (ATS Engineering). The KlBre1 RBD-Rad6 complex was purified by nickel–nitrilotriacetic acid agarose (Ni-NTA, Smart Life sciences) and ion exchange (Hitrap Q HP, GE healthcare) columns, followed by a 2-hour treatment with thrombin (5 units for 1 mg of complex) at room temperature to remove the his-tag on KlRad6, and further purified by gel filtration (Superdex 200 10/300, GE healthcare). Purified complexes were concentrated to 10 mg/ml in a buffer containing 20 mM Tris (pH 7.5), 200 mM sodium chloride and 2 mM dithiothreitol (DTT), flashed cooled in liquid nitrogen and stored at −80 °C.

Unless otherwise indicated, KlBre1 fragment 1-206 containing N-terminal his- and streptavidin-binding (strep-) tags and KlRad6 containing N-terminal his- and hemagglutinin (HA-) tags were used for the biochemical assays. The coding regions for the KlBre1 fragment and the full length KlRad6 were inserted into vector pET28A, and the strep- and HA-tags were introduced by PCR-based mutagenesis. The recombinant double tagged KlBre1 RBD and KlRad6 proteins were expressed in BL21 Rosetta (DE3) cells and purified with Ni-NTA, Heparin (Hitrap Heparin HP, GE healthcare) and gel filtration (Superdex 200 10/300, GE healthcare) columns. The *S. cerevisiae* E1 protein Uba1 and ubiquitin were purified as described(Lee & Schindelin, 2008; Shen *et al*, 2021). Briefly, the gene fragment corresponding to Uba1 residues 10-1024 was inserted into vector pET28A. Uba1 was expressed in BL21 Rosetta (DE3) cells and purified with Ni-NTA, hydrophobic interaction (Hitrap Butyl HP, GE Healthcare) and gel filtration (Superdex 200 Increase 10/300) columns. The ubiquitin gene was inserted into vector pET28A. Ubiquitin was expressed in in BL21 Rosetta (DE3) cells and purified with Ni-NTA, ion-exchange (Hitrap Q HP) and gel filtration (Superdex 200 10/300) columns.

Amino acid substitutions were introduced with a PCR based protocol and verified by DNA sequencing. The substituted proteins were expressed and purified following the same protocols for the wild type proteins.

The selenomethionine (SeMet)-substituted KlRad6 was produced by inhibiting the host methionine production and supplementing SeMet(Doublie *et al*, 1996). The KlBre1 RBD-Rad6 complex containing SeMet-substituted KlRad6 complex was purified following the same protocol for the native complex, except that the DTT concentration is increased to 10 mM in the storage buffer.

### Crystallization and structural determination

Hexagon-shaped crystals of the complex containing the KlBre1 fragment 1-206 and KlRad6 (crystal form 1) were obtained with vapor diffusion sitting drop experiments at 18 °C. The reservoir solution contains 0.2 M sodium acetate (pH 5.5) and 15% PEG3350. Before data collection, the crystals were equilibrated in the reservoir solution supplemented with 25% glycerol, flash cooled and stored in liquid nitrogen. Diffraction data were collected at the Shanghai Synchrotron Radiation Facility (SSRF) Beamline BL19U1 at 0.97853 Å on a Pilatus 6M detector. Diffraction data were indexed, integrated and scaled with the HKL3000 suite(Otwinowski & Minor, 1997). Initial molecular replacement calculations with PHASER(McCoy *et al*, 2007) with the structure of the *Saccharomyces cerevisiae* Rad6 (ScRad6, PDB 1AYZ)(Worthylake *et al*, 1998) as the search model did not yield interpretable electron density maps. An analysis with PAIRCOIL2(McDonnell *et al*, 2006) indicated that the KlBre1 N-terminal region contains at least one coiled-coil of 40 residues in length. A predicted coiled-coil structure was generated with CCFOLD(Guzenko & Strelkov, 2018). Using this structure and the structure of ScRad6 as search models, molecular replacement calculations with PHASER and subsequent density improvement with PHENIX(Adams *et al*, 2010) produced interpretable density maps. Model building was carried out with O(Jones *et al*, 1991) and COOT(Emsley & Cowtan, 2004). Refinement was carried out with PHENIX.

Elongated diamond-shaped crystals of the complex containing the KlBre1 fragment 1-184 and SeMet-substituted KlRad6 (crystal form 2) were obtained with vapor diffusion sitting drop experiments at 18 °C. The reservoir solution contains 0.1 M sodium acetate (pH 4.6) and 1 M ammonium sulfate. Diffraction data was collected on SSRF beamline BL19U1 at 0.97846 Å and processed with the HKL3000 suite. The structure was determined with molecular replacement with PHASER, using the structure of crystal form 1 as the search model. It was refined with PHENIX.

### Pull-down experiments

To probe the KlBre1 RBD-Rad6 interaction, 15 μM his-strep-tagged KlBre1 RBD was incubated with 4 μM KlRad6 in a binding buffer containing 20 mM Tris (pH7.5) and 200 mM sodium chloride on ice for 30 minutes. The reaction mixture was subsequently incubated with 30 μl strep-tactin beads (Smart Lifesciences) equilibrated in the binding buffer for 2 hours at 4°C. After washing the beads twice with the binding buffer, bound proteins were eluted with the binding buffer supplemented with 2 mM desbiotin and analyzed with sodium dodecyl sulphate-polyacrylamide gel electrophoresis (SDS PAGE).

### Isothermal titration calorimetry

Isothermal titration calorimetry (ITC) experiments were performed on a MicroCal PEAQ-ITC instrument (Malvern) at 25 °C. Prior to the ITC experiments, both KlBre1 RBD and KlRad6 were exchanged in a buffer containing 20 mM Tris (pH 7.5) and sodium chloride at indicated concentrations. To characterize binding, a solution containing 100 μM (for experiments with the wild type or Q22A-, K30A-, R35E-, R41E-, R171E-, R179A-substituted KlBre1 RBD with 1 M salt or the wild type KlBre1 RBD with 200 mM salt) or 150 μM (for the experiment with the S111L-subsbituted KlRad6) or 200 μM (for the experiment with the R171E-substitued KlBre1 RBD with 200 mM salt) or 400 μM (for the experiment with the V180A/F181A-substituted KlBre1 RBD) KlRad6 was injected into a 300-μl cell that stores 50 μM KlBre1 RBD, 2 μl at a time. Data were analyzed with ORIGIN 7.0 (Originlab).

### 3-(2-Iodoacetamido)-2,2,5,5-tetramethyl-1-pyrolidinyloxy labeling

For 3-(2-Iodoacetamido)-2,2,5,5-tetramethyl-1-pyrolidinyloxy (IPSL) labeling, 10 μM KlRad6 was incubated with 300 μM IPSL for 15 minutes in a buffer containing 20 mM Tris (pH 7.5) and 200 mM sodium chloride, in the presence or absence of 40 μM KlBre1 RBD. The reaction was stopped by adding 600 μM L-cysteine. The level of IPSL labeling was accessed by mass spectrometry (MS). After tryptic digestion of the reaction mixture, the resulting peptides were separated by nano-liquid chromatography on an easy-nLC 1200 system (Thermo Fisher Scientific) and directly sprayed into a Q-Exactive Plus mass spectrometer (Thermo Fisher Scientific). The MS analysis was carried out in data-dependent mode with an automatic switch between a full MS and a tandem MS (MS/MS) scan in the Orbitrap. For full MS survey scan, the automatic gain control target was set to 1e6, and the scan range was from 350 to 1750 with a resolution of 70,000. The 10 most intense peaks with charge state ⩾ 2 were selected for fragmentation by higher energy collision dissociation with normalized collision energy of 27%. The MS2 spectra were acquired with a resolution of 17,500, and the exclusion window was set at ± 2.2 Da. All MS/MS spectra were searched using the PD search engine (v 1.4.0, Thermo Fisher Scientific) with an overall false discovery rate for peptides less than 1%. Peptide sequences were searched using trypsin specificity allowing a maximum of two missed cleavages. IPSL-Alkylation (+198.137 Da) on cysteine, acetylationon peptide N-terminal and oxidation of methionine were specified as variable modifications. Mass tolerances for precursor ions were set at ± 10 ppm for precursor ions and ± 0.02 Da for MS/MS. The ratio of labeled peptides to unlabeled peptides in the mass spectrometric analysis was calculated to represent the level of IPSL labeling.

### Free ubiquitin chain formation

The reaction mixture for the free ubiquitin chain formation experiments contains 60 mM Tris (pH 8.5), 50 mM sodium chloride, 50 mM potassium chloride, 10 mM magnesium chloride, 0.1 mM DTT, 3 mM ATP, 90 nM Uba1, 60 μM ubiquitin and 10 μM (for experiments presented in Figs 3B and S7B-C) or 5 μM KlRad6 (for experiments presented in Figs 3C and S7A). When indicated, 20 μM KlBre1 RBD were supplemented. After incubating at 30 °C for the indicated period, the reactions were stopped by boiling in the SDS PAGE loading buffer. To remove KlBre1 RBD, the boiled reaction mixture was subject to strep-tactin beads precipitation twice and the supernatant was collected for analysis. For comparison, reactions presented in figures 3B-C without KlBre1 RBD were supplemented with the same amount of KlBre1 RBD after the reaction and subjected to the same treatment with strep-tactin beads. The reaction mixtures were analyzed with SDS PAGE and western blot with an anti-ubiquitin antibody (sc-8017, Santa Cruz Biotechnology, RRID: AB_628423, 1:2000 diluted). To probe ubiquitin chains attached to the KlBre1 RBD during the reaction, reaction mixtures prior to the treatment with strep-tactin beads were analyzed with western blot with an anti-strep antibody (AE066, ABclonal, RRID: AB_2863792, 1:5000 diluted).

### Rad6~ubiquitin discharging

For the discharging reactions, the N-terminal his-tag in KlRad6 and ubiquitin were removed. After incubating the proteins with thrombin (5 units for 1 mg of protein) at room temperature for 2 hours, the untagged proteins were purified with gel filtration (Superdex 200 increase 10/300). When indicated, KlBre1 RBD (1-184) was supplemented to discharging reactions with the wild type KlRad6 (Fig. S7E). The corresponding KlBre1 gene fragment was inserted into vector pET28A. The recombinant protein contains an N-terminal his-tag and was purified by Ni-NTA, Heparin (Hitrap Heparin HP) and gel filtration columns (Superdex 200 10/300). For discharging reactions with the wild type KlRad6 (Fig. S7E), KlRad6 was first charged with ubiquitin with a reaction mixture containing 60 mM Tris (pH 8.0), 50 mM sodium chloride, 50 mM potassium chloride, 10 mM magnesium chloride, 0.1 mM DTT, 3 mM ATP, 30 nM Uba1, 50 μM ubiquitin and 2 μM KlRad6. After a 10-minute incubation at 30 °C, the charging reaction was stopped by adding 50 mM EDTA (pH 8.0) and ubiquitin discharge from the KlRad6~ubiqtuin conjugate in the presence or absence of 20 μM KlBre1 RBD (1-184) were allowed to proceed at 30 °C. After the indicated period, aliquots of the reaction were removed and analyzed by non-reducing SDS-PAGE. For the discharging reactions with the S111L-substituted KlRad6 (Figs 3D and S7F), S111L-substituted KlRad6 was first charged with I44A/K0-substituted ubiquitin with a reaction mixture containing 60 mM Tris (pH 8.0), 50 mM sodium chloride, 50 mM potassium chloride, 10 mM magnesium chloride, 0.1 mM DTT, 3 mM ATP, 300 nM Uba1, 20 μM I44A/K0-substituted ubiquitin and 10 μM S111L-substituted KlRad6. The reaction mixture was incubated at 30 °C for 30 minutes, with additional 300 nM Uba1 and 3 mM ATP supplemented at minute 10 and 20. The charging reaction was stopped by adding 50 mM EDTA (pH 8.0) and the discharging reaction was initiated by supplementing 60 μM of I44A-substituted ubiquitin to the reaction mixture. When indicated, 30 μM of wild type or substituted KlBre1 RBD (1-206) was also supplemented. The reaction mixture was incubated at 30 °C for the indicated period and analyzed by non-reducing SDS PAGE followed by western blot with an anti-ubiquitin antibody (sc-8017, Santa Cruz Biotechnology, RRID: AB_628423, 1:2000 diluted). For quantification, band intensities were read by ImageJ (https://imagej.nih.gov/ij/).

### Yeast strains and plasmids

Yeast strains used are listed in Supplemental Table 1. All strains used in this study are derivates of JKM139 (*ho MAT**a** hml::ADE1 hmr::ADE1 ade1-100 leu2-3,112 trp1::hisG’ lys5 ura3-52 ade3::GAL::HO*) or tGI354 (*MATa-inc arg5,6::MATa-HPH ade3::GAL::HO hmr::ADE1 hml::ADE1 ura3-52*) Strains were constructed with standard yeast genetic manipulation. Mutant strains were confirmed by PCR or sequencing.

### Replication stress/DNA damaging reagents sensitivity test

Sensitivity towards replication stress/DNA damaging reagents was tested using a spotting assay. A serial dilution of overnight yeast cultures were produced and 5μl aliquots of the diluted culture were spotted onto YPD plates with the indicated replication stress/DNA damaging agents. Plates were incubated at 30 °C for 3-4 days before analysis.

### Analysis of DSB repair by ectopic recombination

To assess the cell survival due to successful DSB repair by ectopic recombination, cells were cultured in YEP-Raffinose to the log phase and subsequently diluted and plated on YEPD or YEP-galactose plates, on which the HO expression is induced. Cells were allowed to grow at 30°C for 3 to 5 days. Survival rate is defined as the number of colonies grown on YEP-galactose plates divided by the number of colonies grown on YEPD plates times the fold of dilution. Five independent experiments were performed for each strain.

### Cellular H2Bub1 level analysis

Whole-cell yeast extracts were prepared using a trichloroacetic acid method as previously described(Chen *et al*, 2012). Samples were resolved with 12% SDS-PAGE and transferred onto a PVDF membrane (Immobilon-P, Millipore). Flag-tagged H2B was detected with western blot using mouse anti-FLAG (F3165, Sigma-Aldrich, RRID: AB_259529, 1:3000 diluted) and HRP-conjugated anti-mouse (sc-516102, Santa Cruz Biotechnology, RRID: AB_2687626, 1:6000 diluted) antibodies.

## Data availability

Diffraction data and refined structures of crystal forms 1 and 2 of the KlBre1 RBD-Rad6 complex have been deposited into the protein data bank (www.rcsb.org), with accession codes 7W75 and 7W76, respectively.

## Acknowledgements

We thank scientists at the National Facility for Protein Science beamline BL19U1 at Shanghai Synchrotron Radiation Facility for setting up the beamline and assistance during diffraction data collection, the Large Equipment Sharing Platform at Tianjin Medical University for assistance with ITC and MS experiments, Drs Hang Zhang and Miaomiao Shen at Tianjin Medical University for their contribution in the early phases of the project.

## Supplementary figures and tables

**Figure S1.**
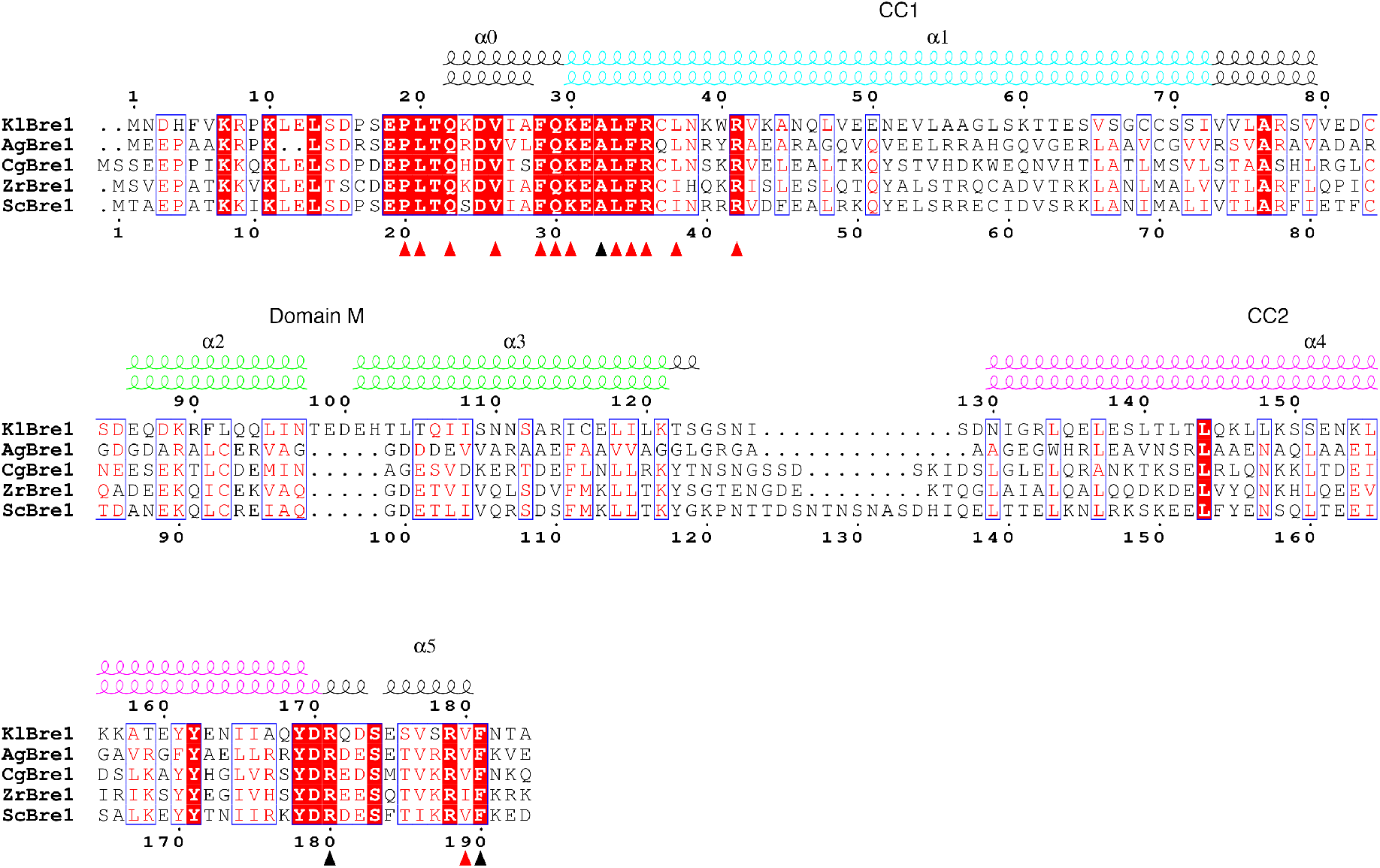
Sequence alignment of the N-terminal region in fungal Bre1 proteins. Residues numbers for KlBre1 and ScBre1 are indicated above and below the alignment, respectively. Secondary structure elements for the two polypeptides in KlBre1 RBD are indicated. The red triangles indicate residues mediating interactions with KlRad6 and between the two KlBre1 polypeptides in the RBD. The black triangles indicate residues mediating interactions with KlRad6 only. AgBre1, *Ashbya gossypii* Bre1; CgBre1, *Candida glabrata* Bre1; ZrBre1, *Zygosaccharomyces rouxii* Bre1.

**Figure S2.**
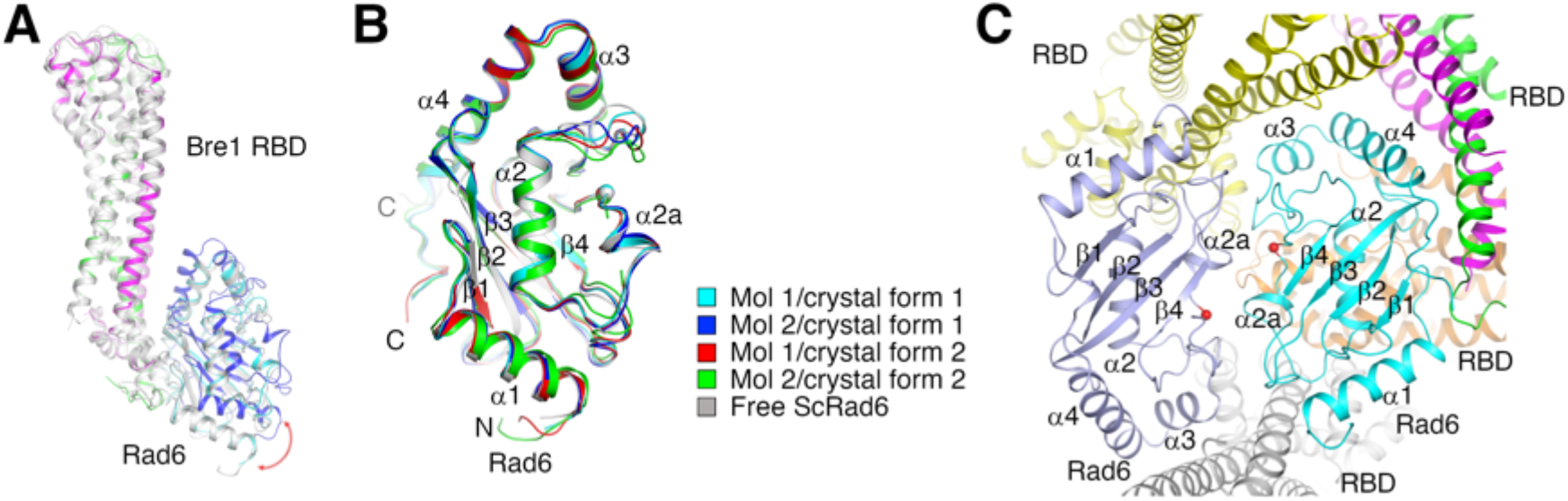
Structure of the KlBre1 RBD-Rad6 complexes in the crystal. (A) Structural alignment of the 4 complexes found in crystal forms 1 and 2. The KlBre1 RBD in these complexes are aligned. The pairwise root mean square deviations (RMSDs) for the equivalent Ca atoms in these KlBre1 RBD molecules are between 0.77 Å and 1.39 Å. Complex 1 in crystal form 1 is colored as in Fig. 1A, KlRad6 molecule 1 in crystal form 2 is colored in blue, other molecules are colored in gray. The red arrow indicates the large structural difference between KlRad6 molecule 1 in crystal form 2 and other KlRad6 molecules in the crystal. (B) Structural alignment of the four KlRad6 molecules found in crystal forms 1 and 2. Structure of the free ScRad6 (PDB 1AYZ)(Worthylake *et al*, 1998) is aligned for comparison. The pairwise RMSDs for the equivalent Ca atoms in the four KlRad6 molecules and the free ScRad6 molecule are between 0.62 Å and 1.16 Å. The spheres indicate Rad6’s active site. (C) KlRad6 molecule 1 in crystal form 2 mediate extensive crystal packing interactions. The KlBre1 RBD-Rad6 complex 1 in crystal form 2 is colored as in Fig. 1a. Protein molecules interacting with the KlRad6 molecule in this complex are highlighted with different colors. The red spheres indicate KlRad6’s active site. RBD, KlBre1 RBD. Structural elements surrounding the active site of this KlRad6 molecule interact with the same region in its symmetry equivalent, burying 1300 Å^2^ of surface area at the interface. In addition to the KlBre1 RBD dimer it forms a complex with, this KlRad6 molecule also interacts with three KlBre1 RBD dimers in the crystal, the related interfaces bury 880, 680 and 580 Å^2^ of surface area.

**Figure S3.**
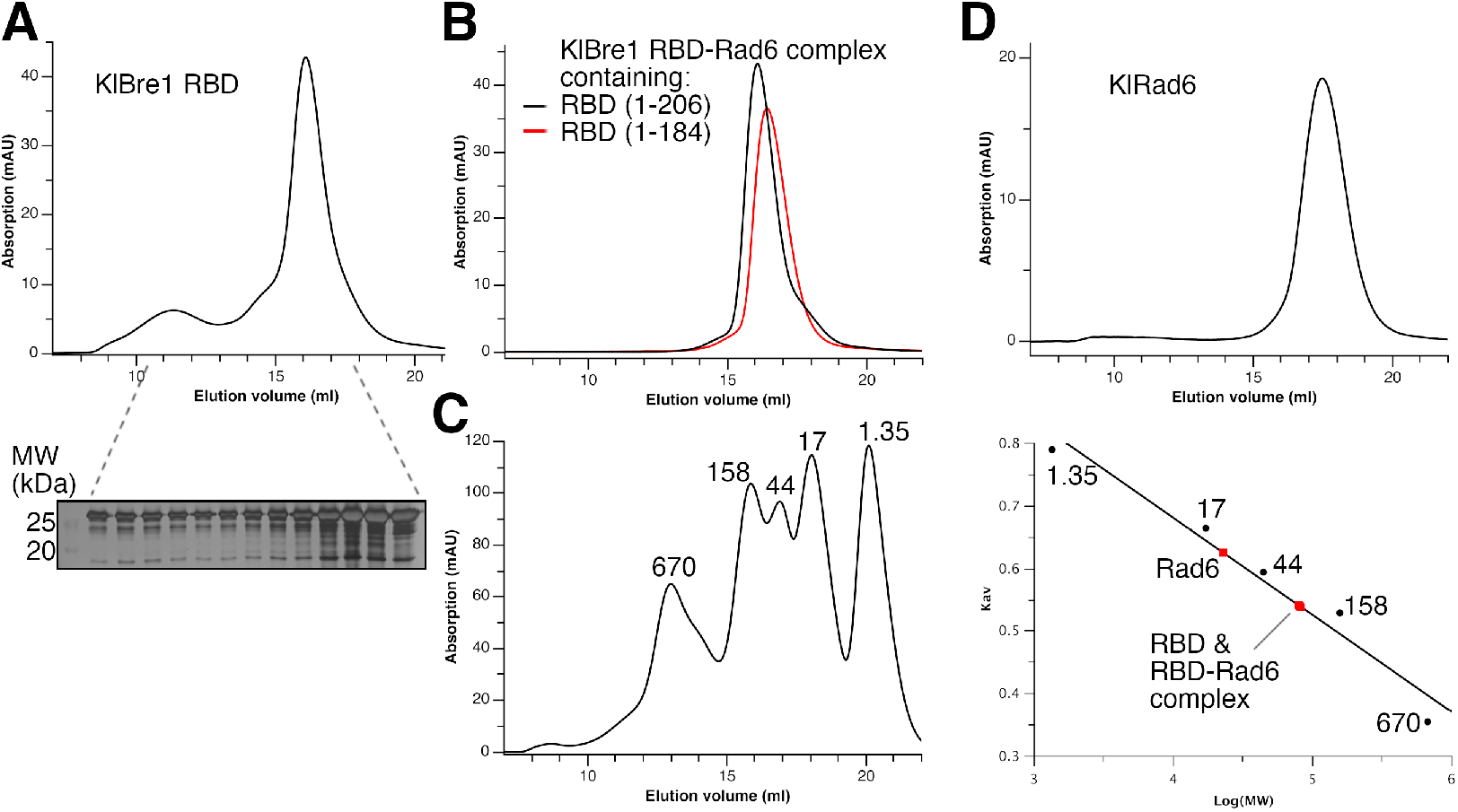
Gel filtration characterization of KlBre1 RBD, KlRad6 and their complexes. (A) Gel filtration characterization of KlBre1 RBD. The lower panel shows a SDS PAGE analysis of the gel filtration experiment. Source data is provided in figure S3 source data 1. (B) Gel filtration characterization of KlBre1 RBD-Rad6 complexes used for crystallization. (C) Calibration of the gel filtration column. The column is calibrated with thyroglobulin, γ-globulin, ovalbumin, myoglobin and vitamin B12 (BIO-RAD, their molecule weights are indicated in units of kDa in the figure). The left and right panels show the elution profile of a mixture of these molecules and the calibration curve, respectively. In the right panel, Kav and the logarithm of the molecular weight of these molecules are fitted to a linear equation. Kav is defined as (Ve-Vo)/(Vt-Vo) (Ve, elution volume; Vo, column void volume (7.2 ml); Vt, total column volume (23.6 ml)). Molecular weight for KlBre1 RBD, KlRad6 and their complexes deduced from the calibration curve are indicated with the red dots. (D) Gel filtration characterization of KlRad6. The gel filtration experiments were performed with a Superdex 200 10/300 column (GE healthcare) with a buffer containing 20 mM Tris (pH 7.5) and 200 mM sodium chloride.

**Figure S4.**
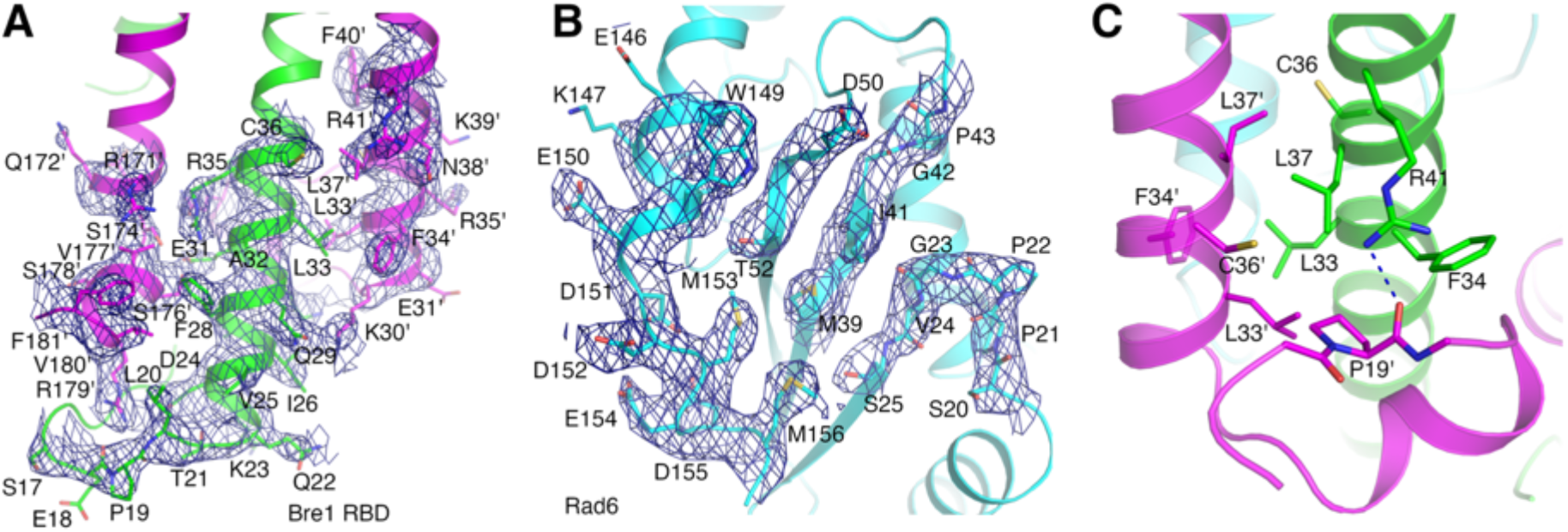
Residues at the KlBre1 RBD-Rad6 interface. (A) and (B) Electron density map at the KlBre1 RBD-Rad6 interface in KlBre1 RBD (A) and KlRad6 (B). Refined electron density map for residues in complex 1/crystal form 1 at the interface and surrounding area is shown in blue. The map is contoured at 1 σ. (C) Interactions between polypeptides 1 and 2 in KlBre1 RBD mediated by residues also participate in the KlBre1 RBD-Rad6 interaction.

**Figure S5.**
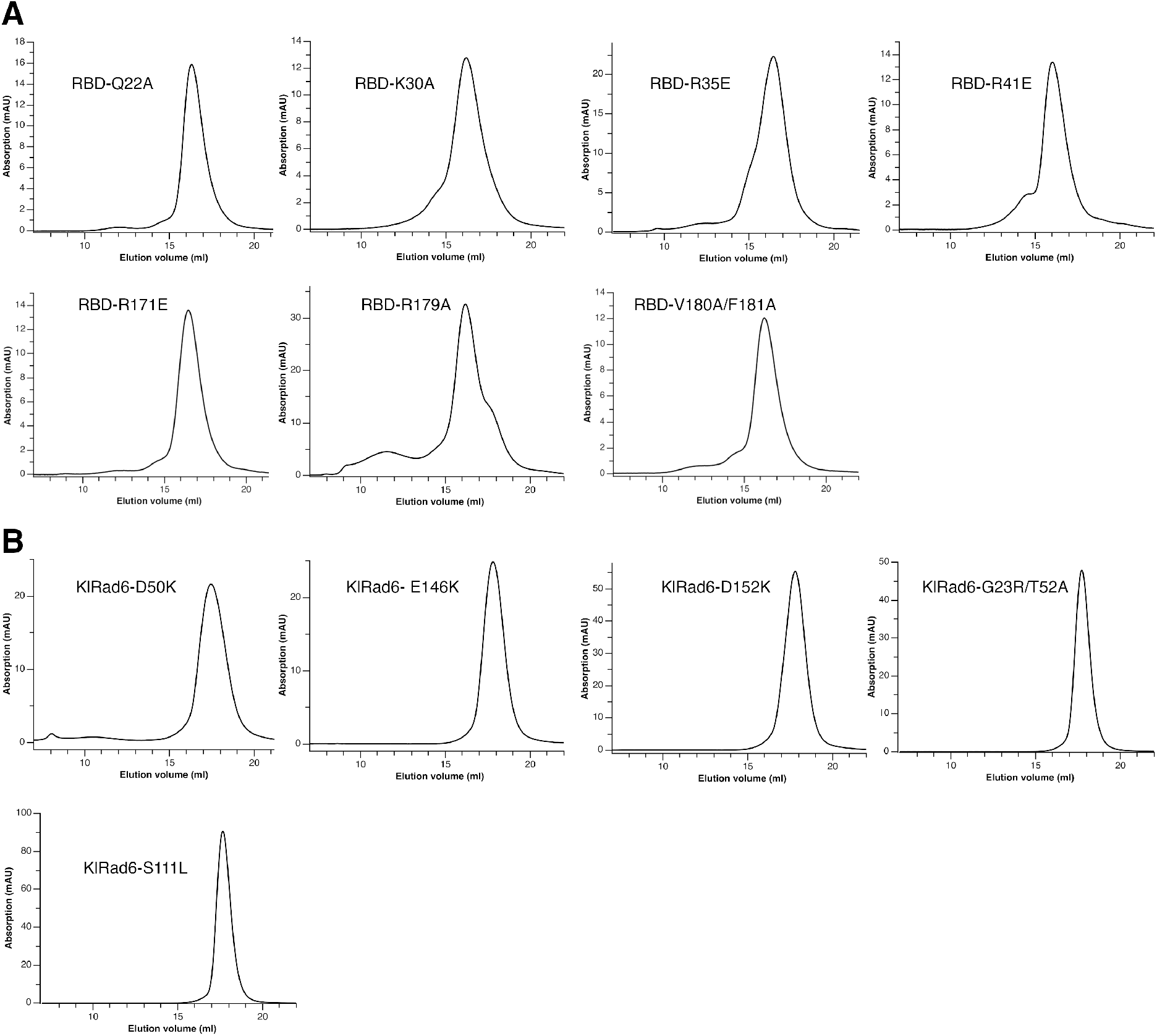
Gel filtration characterization of substituted KlBre1 RBD (A) and KlRad6 (B). The gel filtration experiments were performed with a Superdex 200 10/300 column (GE healthcare) with a buffer containing 20 mM Tris (pH 7.5) and 200 mM sodium chloride.

**Figure S6.**
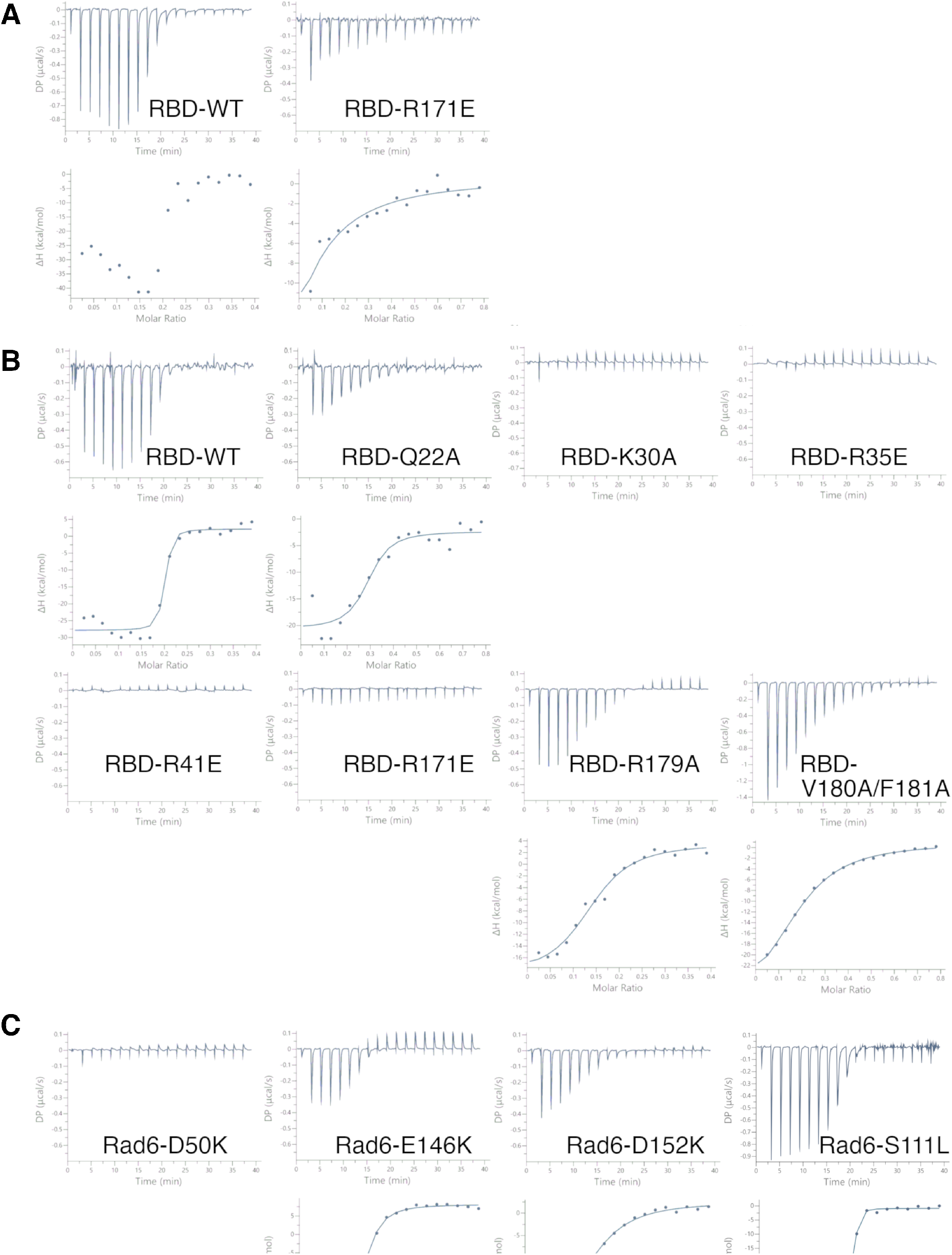
ITC experiments. (A) ITC experiments with 200 mM salt and the wild type KlRad6. (B) ITC experiments with 1M salt and the wild type KlRad6. (C) ITC experiments with 1M salt and the wild type KlBre1 RBD.

**Figure S7.**
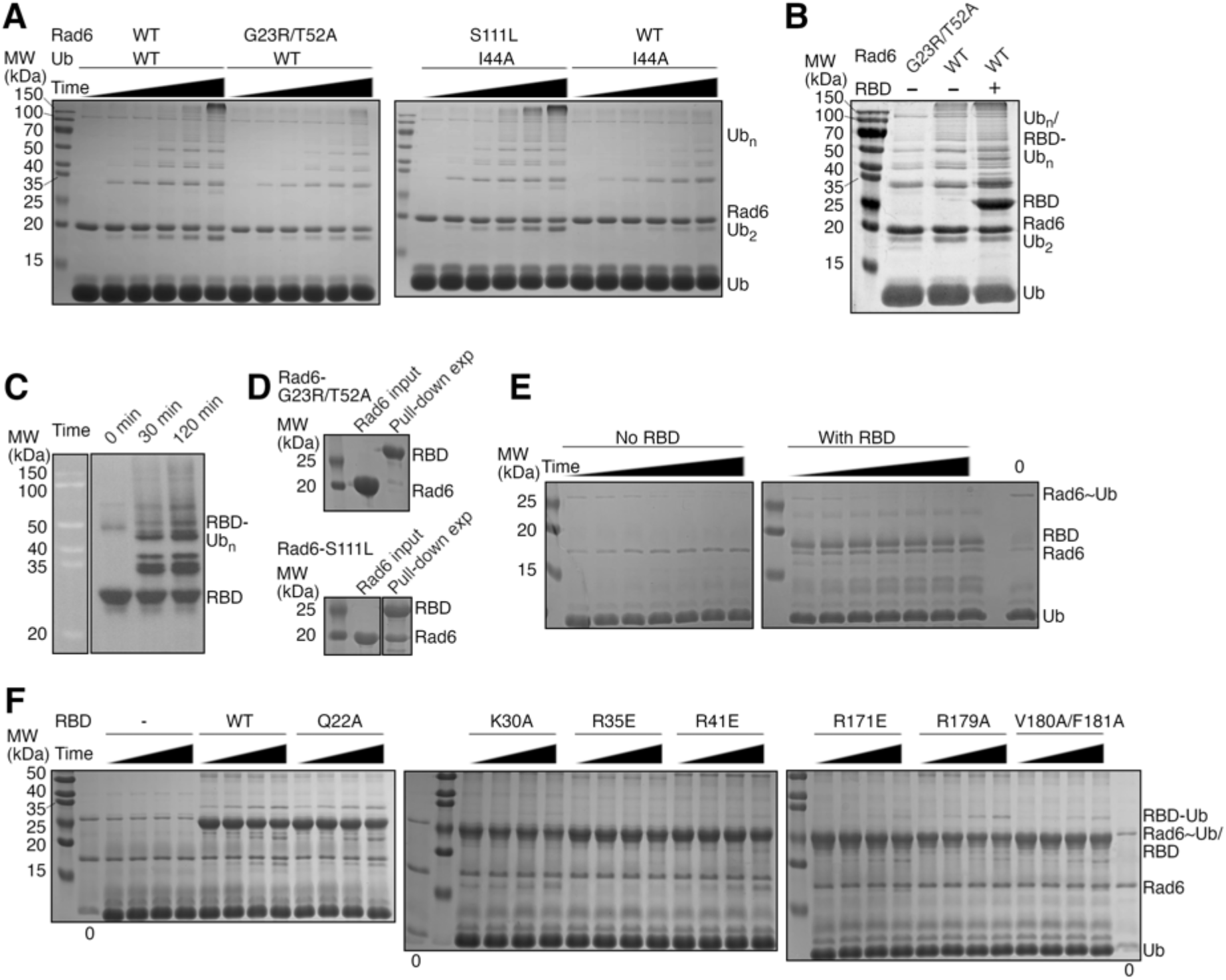
Rad6 activity assays. (A) and (B) Ubiquitin chain production by KlRad6. SDS PAGE analysis of the reactions are presented. Experiments presented in panel A were carried out in the absence of KlBre1 RBD for 0, 5, 10, 20, 30 and 60 minutes. Reactions presented in panel B were carried out for 30 minutes. KlBre1 RBD was not removed before analysis. Ub, ubiquitin; Ub2 and Ubn, ubiquitin chains with 2 or n ubiquitin moieties, respectively. (C) KlBre1 RBD can be ubiquitinated. Western blot analysis against the streptavidin-binding peptide fused to KlBre1 RBD N-terminus after the ubiquitin chain formation reaction is presented. The reactions were allowed to proceed for the indicated time. (D) Pull-down experiments probing the KlBre1 RBD-Rad6 interaction. Experiments with the G23R/T52A- and S111L-substituted KlRad6 are presented. (E) and (F) Non-reducing SDS PAGE analysis of the KlRad6~ubiquitin discharging reaction. Lanes marked with “0” shows SDS PAGE analysis of the KlRad6~ubquitin conjugate prior to the reaction. In reactions presented in panel E, wild type ubiquitin, KlRad6 and KlBre1 RBD (1-184) were used. The reactions were allowed to proceed for 10, 20, 40, 70, 100, 130 and 150 minutes before analysis. In reactions presented in panel F, K0/I44A-substituted ubiquitin was charged to S111L-substituted KlRad6 and discharged to I44A-substituted ubiquitin in the absence or presence of wild type or substituted KlBre1 RBD (1-206). The reactions were allowed to proceed for 5, 10, 20 and 40 minutes before analysis. Source data for panels A-F are provided in figure S7 source data 1.

**Figure S8.**
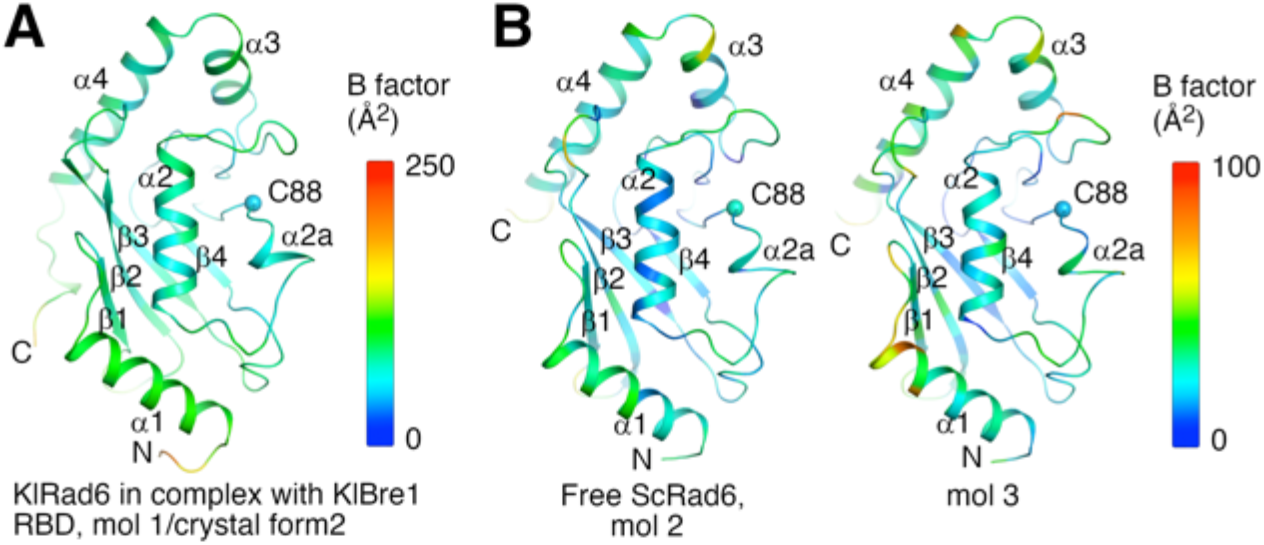
Temperature factor distribution of Rad6 molecules in crystal structures. (A) Temperature factor distribution of KlRad6 molecule 1 in crystal form 2 in our study. (B) Temperature factor distribution of molecules 2 and 3 in the crystal of free ScRad6. The active site (C88) of the Rad6 molecules are indicated.

**Figure S9.**
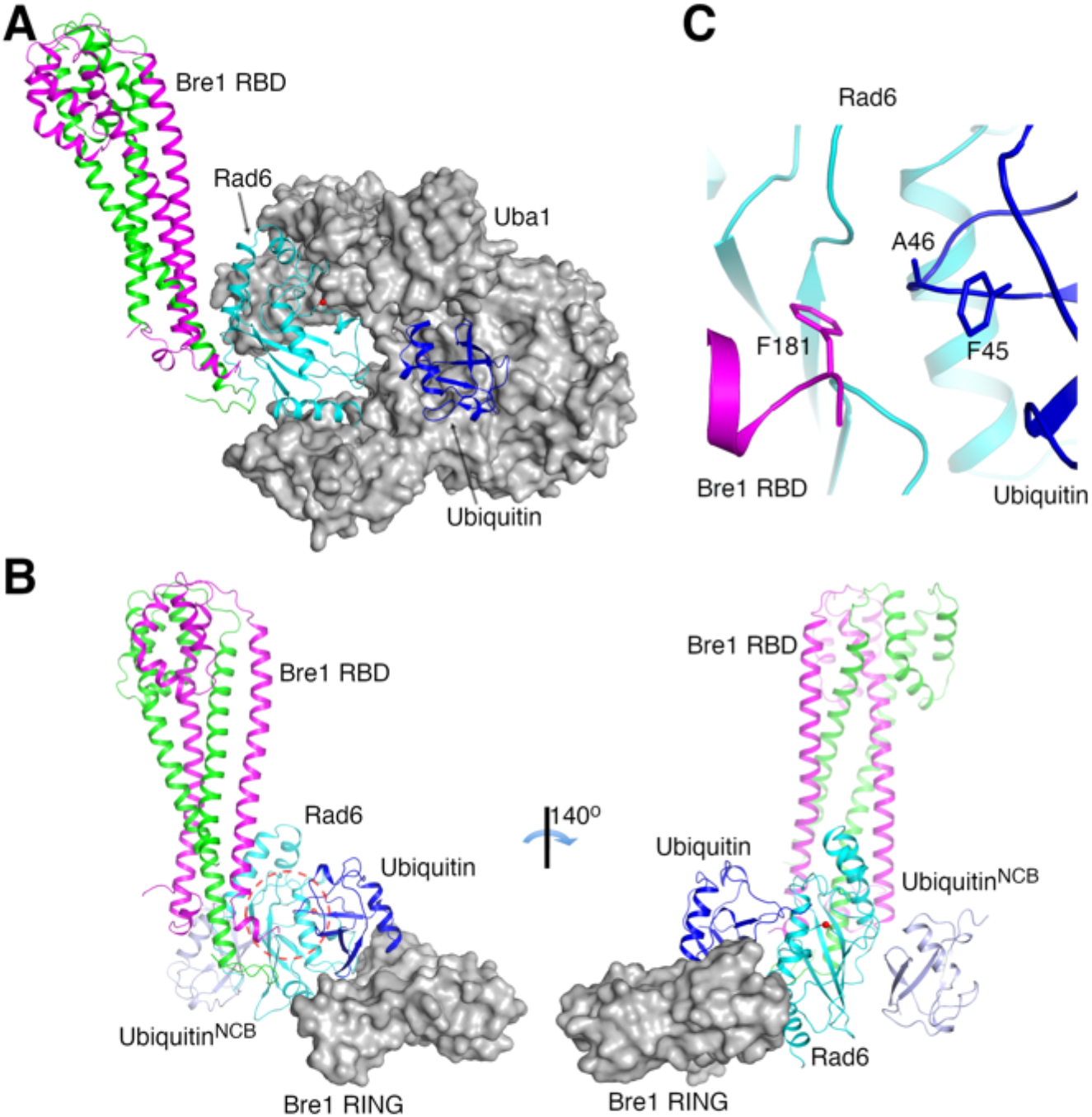
Structural modeling of Bre1 RBD during the H2Bub1 catalysis. (A) Model of the Bre1 RBD-Rad6-E1-ubiquitin complex in the ubiquitin charging reaction. The E1 enzyme (Uba1) and ubiquitin are modeled based on the structure of the Uba1-Ubc4-ubiquitin complex (PDB 4II2)(Olsen & Lima, 2013). (B) Model of the Bre1 RBD-Rad6-ubiquitin-Bre1 RING domain complex in the ubiquitin discharging reaction. Bre1’s RING domain and ubiquitin are modeled based on the structure of the RNF4 RING domain in complex with UbeV2, the ubiquitin-Ubc13 complex in the closed conformation (PDB 5AIT)(Branigan *et al*, 2015) and the structure of the Bre1 RING domain (PDB 4R7E)(Kumar & Wolberger, 2015). The ubiquitin molecule bound to the non-canonical back side of Rad6 (ubiquitin^NCB^)(Kumar *et al*, 2015) is shown. (C) Potential interaction between Bre1 RBD and ubiquitin stabilizing the closed conformation of the bound Rad6~ubiqutin conjugate. The region marked with the red circle in panel B is shown, important residue side chains are highlighted. Uba1 and the Bre1 RING domain are presented in surface presentation in gray. The red spheres indicate Rad6’s active site.

**Figure S10.**
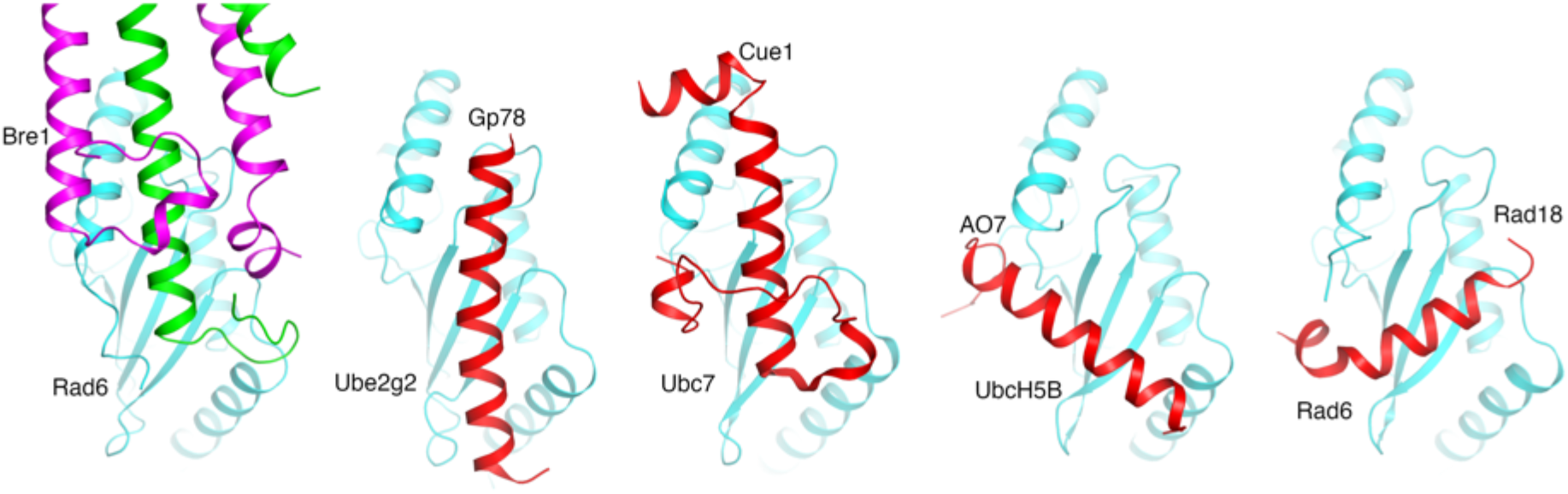
E3 enzymes bind to the back side of E2 enzymes with drastically different mechanisms. Structures of the KlBre1 RBD-Rad6 complex and the E2BBRs in Gp78 (PDB 3H8K)(Das *et al*, 2009), Cue1 (PDB 4JQU)(Metzger *et al*, 2013), AO7 (PDB 5D1K)(Li *et al*, 2015) and Rad18 (PDB 2YBF)(Hibbert *et al*, 2011) in complex with the related E2 enzymes are shown. The E2 enzymes are aligned.

**Table S1.** Yeast strains

| Strain name | Parental strain | Genotype | Source |
| --- | --- | --- | --- |
| JKM139 |  | *MATa ho hml::ADE1 hmr::ADE1 ade1-100 leu2-3,112 TRP1::hisG lys5 ura3-52 ade3::GAL::HO* | Lee et al (Lee et al, 1998) |
| tGI354 |  | *MATa-inc arg5,6::MATa-HPH,ade3::GAL::HO hmr::ADE1 hml::ADE1 ura3-52* | Ira et al (Ira et al, 2003) |
| yZSH241 | tGI354 | *bre1::TRP1* | this study |
| yMF001 | tGI354 | *bre1::TRP1 + pRS316-bre1-Q23A* | this study |
| yMF002 | tGI354 | *bre1::TRP1 + pRS316-bre1-Q30AK31D* | this study |
| yMF003 | tGI354 | *bre1::TRP1 + pRS316-bre1-R36ER42E* | this study |
| yMF004 | tGI354 | *bre1::TRP1 + pRS316* | this study |
| yMF005 | tGI354 | *bre1::TRP1 + pRS316-BRE1* | this study |
| yMF006 | JKM139 | *bre1::TRP1 + pRS316-bre1-Q23A* | this study |
| yMF007 | JKM139 | *bre1::TRP1 + pRS316-bre1-Q30AK31D* | this study |
| yMF008 | JKM139 | *bre1::TRP1 + pRS316-bre1-R36ER42E* | this study |
| yMF009 | JKM139 | *bre1::TRP1 + pRS316* | this study |
| yMF010 | JKM139 | *bre1::TRP1 + pRS316-BRE1* | this study |
| yLD002 | JKM139 | *bre1::TRP1* | this study |
| yMF011 | JKM139 | *KanMX-FLAG-H2B bre1::TRP1 + pRS316-bre1-Q23A* | this study |
| yMF012 | JKM139 | *KanMX-FLAG-H2B bre1::TRP1 + pRS316-bre1-Q30AK31D* | this study |
| yMF013 | JKM139 | *KanMX-FLAG-H2B bre1::TRP1 + pRS316-bre1-R36ER42E* | this study |
| yMF014 | JKM139 | *KanMX-FLAG-H2B bre1::TRP1 + pRS316* | this study |
| yMF015 | JKM139 | *KanMX-FLAG-H2B bre1::TRP1 + pRS316-BRE1* | this study |
| yGX060 | JKM139 | *KanMX-FLAG-H2B bre1::TRP1* | this study |

